# Detection and Classification of Cervical Cancer Using Optimized Deep Learning Approach

**DOI:** 10.1101/2025.06.05.657998

**Authors:** S Lalitha, B V Baiju, Sandeep Kumar Mathivanan, Partheeban Nagappan, Chandra Shakher Tyagi, Saurav Mallik

## Abstract

Given the global impact of the cervical cancer epidemic on people’s health, it is critical to have readily available and effective screening technologies. In order to effectively fight this disease, it is crucial to identify the groups who are most at risk. Our study aims to build a robust deep learning system tailored for the classification of cervical cancer using images acquired from Pap screenings; this will allow us to tackle this challenge head-on. Our approach improves upon previous methods of visual feature detection by using a deep learning model based on transfer learning called Squeeze-and-Excitation-ResNet152. The Deer Hunting Optimization method is used to optimise the network by modifying its hyper-parameters. We test our methods on eleven distinct disease sets, with a total of 8838 images distributed differently. Sources such as CRIC and SIPaKMeD were consulted for the acquisition of these images. In order to reduce dataset bias, we use a cost-sensitive loss function all through training. Impressive performance metrics were generated by the testing set analysis, which significantly outperformed the previous methods. Accuracy was 99.74%, precision was 98.98%, recall was 98.15%, specificity was 98.97%, and F1-Score was 99.06%. We can greatly enhance the identification of problems connected to cervical cancer with our technologies. One way to achieve this goal is to make it easier for medical professionals to make quick diagnosis.

## 1. Introduction

With the regrettable distinction of being the fourth most frequent cancer among women worldwide and the seventh most common cancer overall, cervical cancer is an important global health problem. The aforementioned disease is closely associated with a number of factors, such as early sexual initiation, inadequate menstrual hygiene practices, the use of oral contraceptives, smoking habits, weakened immune systems, early pregnancy, and multiple sexual partners [1]. All of these things put you at a higher chance of getting cervical cancer, which highlights the need of early identification and complete prevention. The traditional approach to cervical cancer diagnosis usually involves cytopathology screening. During a cervical cytopathological examination, medical personnel gently remove cells from a patient’s cervix’s surface using specialised brushes [2]. After being gathered, these cells are carefully put onto a glass plate for further examination. After that, expert cytopathologists carefully examine these removed cells under a microscope in an effort to spot any anomalies or signs of cancer. But because each slide may have a large number of cells on it, manual examination becomes quite complex and prone to human mistake, which presents difficulties for experts [3]. As a result, there is a pressing need to create and use more effective and dependable remedies to lessen these problems with cervical cancer diagnosis. The creation of automated computer-aided diagnostic (CAD) systems seems to be the answer to this problem. These devices represent a major advancement in the diagnosis of cervical cancer by enabling the quick and reliable examination of Pap slides [4]. At first, classical machine learning and image processing methods served as the basis for CAD systems. However, the use of manually created characteristics sometimes led to less-than-ideal classification results, which reduced the precision of diagnosis. Nevertheless, CAD systems have undergone a revolution because to recent developments in deep learning technology, which have removed earlier constraints and greatly enhanced their performance in a number of criteria [5]. CAD systems have improved their ability to detect abnormalities in Pap slides by using deep learning algorithms. This has allowed for more accurate and early diagnosis of cervical cancer. Technological developments are a major source of innovation and growth in many sectors, including computer vision, medical imaging, natural language processing (NLP), and more [6]. Deep learning has become a ground-breaking paradigm in the field of machine learning in this context. With this advanced method, computers are able to recognise complex patterns and extract relevant characteristics on their own. They are especially good at performing classification jobs with a high degree of accuracy and productivity [7].

Nonetheless, the availability of extensive labelled training datasets is a critical component in the efficacy of deep learning models. The foundation for the model’s development of its comprehension of the underlying patterns and connections in the data it encounters is provided by these datasets. Thus, obtaining and maintaining large-scale labelled datasets are essential preconditions for the effective implementation of deep learning solutions in many fields. Convolutional Neural Networks (ConvNets) are a fundamental architecture in the field of deep learning [8]. ConvNets, which are highly regarded for their abilities to analyse visual data, are being used for many different tasks, such as segmentation, classification, and image recognition. They perform accurately and robustly in a variety of visual identification tasks because to their hierarchical structure, which makes it possible for them to effectively record complex characteristics and spatial hierarchies within images. This work focuses on the complex domain of heterogeneous cervical cancer cases, which poses significant challenges to categorization when compared to conventional approaches [9].

The Human Papilloma Virus (HPV) is a significant cause of cancer-related death, with 570,000 women worldwide affected by cervical cancer. The problem is made worse by the lack of oncologists and the limited resources in poor countries. Researchers have created CervixNet, a deep learning approach for the detection and classification of cervical cancer, in order to overcome these difficulties. CervixNet uses a Hierarchical Convolutional Mixture of Experts (HCME) method to improve the quality of cervigram images and segment the Region of Interest (RoI). Using publicly accessible datasets from Intel and Mobile-ODT Kaggle, the technique gets a kappa score of 0.951 and an accuracy of 96.77%, indicating its potential as an affordable option for first cervical cancer screening [10].

Predicting life-threatening conditions like cancer, renal failure, and heart attacks is becoming more and more dependent on machine learning and deep learning. This paper suggests a very advanced method for using ML systems to predict cervical cancer. Four stages comprise the research methodology: preprocessing, choosing predictive models, collecting datasets, and delineating algorithmic procedures. Decision trees, logistic regression, support vector machines, adaptive boosting, gradient boosting, random forest, and XGBoost were among the many machine learning algorithms that were used. According to the studies, support vector machines scored an astounding 99% accuracy, while random forest, decision tree, adaptive boosting, and gradient boosting algorithms earned the maximum classification score of 100%. These results point to the possibility of using machine learning (ML) techniques to enhance cervical cancer early detection and intervention programmes [11].

In order to tackle these issues, a thorough inquiry is started, starting with a well-planned experiment that seeks to determine the best ConvNet model. After this first stage, the study explores a thorough assessment of several performance enhancement methods. This means using state-of-the-art image improvement techniques and carefully examining optimizer selection procedures. The research attempts to improve and enhance the performance of the suggested categorization framework via this iterative procedure. The integration of deep learning techniques’ high performance and strong generalisation capabilities is a fundamental component of the suggested methodology. With the help of deep learning algorithms’ flexibility and adaptability, the framework shows that it can handle the challenges posed by cervical cancer categorization jobs. One of the most important recommendations that come out of this study is to use transfer learning methods in a hybrid deep learning network model. Transfer learning enables the smooth transfer of learnt characteristics, enabling the model to attain increased efficiency and accuracy in the categorization of various forms of cervical cancer problems. This is achieved by using the richness of information extracted from pre-existing datasets and assignments. This deliberate use of transfer learning speeds up the learning process and improves the model’s capacity to identify minute differences and subtlety in Pap smear images, which in turn allows for more precise and clinically meaningful diagnosis of cervical cancer variants. The contribution of this research work is,

- The image pre-processing methods enhance the quality of image data by minimizing unwanted distortions that could obscure important features. This improvement is achieved systematically through mathematical morphological procedures, aimed at refining image characteristics to boost classification accuracy.
- Following pre-processing, an optimized SQ-ResNet152 model, employing transfer learning, is used for the classification task. This advanced model excels at recognizing patterns and features in the processed image data, effectively classifying Pap smear images into 11 categories, leveraging the expansive CRIC and SIPaKMeD datasets.
- To ensure robust classification outcomes, the proposed network model adeptly extracts reliable features from medical images. Through the application of DHO optimization techniques, ConvNet hyperparameters are fine-tuned, significantly enhancing diagnostic accuracy for various illnesses.
- The proposed method is evaluated on publicly available datasets using the Python platform. Experimental results indicate that the approach outperforms all other contemporary methods in terms of efficiency.

The structure of the article has been assessed so as to meet the learning objectives that have been preset. This Section 2 is to assess the clinical effectiveness of the latest advanced Pap smear categorization methods. Further, additional deep learning-based architectures aimed at the classification of Pap smear images have been explored in Section 3. This section covers such aspects as design, training cycles, and performance parameters of such complex models. In Section 4, details related to the experimental design, evaluation criteria, and key findings of the empirical study are provided. This section provides the empirical basis for the study by presenting the data sets, experimental methods and evaluation criteria used for the proposed models. This section offers an overview of these options and evaluates their merits and limitations. This information and skills drawing synthesis are performed in Section 5 to conclude the essay with all discussed previously parts. In addition, this section proposes areas of further work and ideas for future improvement of deep learning approaches to Pap smear image analysis. Present within the scope of the current study is Section 5 which presents the future directions that would make positive contributions towards cervical cancer diagnosis and treatment.

## 2. Related Work

Shenghua Cheng [12] presented a unique method for lesion cell identification by combining high- and low-resolution whole slide images (WSIs) for lesion cell identification. A large dataset of 3,545 patient-level WSIs, including 79,911 annotations from various medical institutions and imaging devices, is used to verify the technique. Impressive performance numbers are achieved by the system, such as an 88.5% true positive rate for identifying the top 10 lesion cells on 447 positive images, and a Specificity of 93.5% and Sensitivity of 95.1% for slide classification. The technology improves the accuracy, efficiency, and scalability of cervical cancer screening by analysing a single giga-pixel WSI in 1.5 minutes.

Hua Chen [13] an AI system called CytoBrain was created to quickly identify aberrant cervical cells, which would simplify clinical diagnosis. Its three main parts are a segmentation module that uses the CompactVGG network to extract cell images from entire slide images, a classification module that uses the same network, and a visualised module for human-aided diagnosis. CytoBrain is an effective tool for widespread cervical cancer screening programmes since it performs better than VGG11 in terms of classification accuracy and time efficiency.

Hiam Alquran [14] in order to facilitate a computer-aided identification of problems related to cervical cells, this research offers a technique that divides pap smear images into seven categories. To distinguish between the seven classes, the system makes use of a Support Vector Machine classifier and automated features that were retrieved using ResNet101. When identifying normal instances and differentiating between normal and abnormal cases, the system reaches 100% accuracy and sensitivity. Additionally, it accurately detects and classifies high-level abnormalities with high accuracy, and it diagnoses instances of mild and moderate dysplasia with an accuracy of around 92%. Five polynomial SVM classifiers make up the cascade structure of the system, which achieves an overall accuracy of 100% during training and 97.3% during testing. This approach may increase women’s survival rates by facilitating early cervical cancer detection.

Lidiya Wubshet [15] the fourth most common disease in the world among women, cervical cancer is becoming more common both in incidence and death, particularly in emerging nations. Conventional screening techniques such as histopathological investigations, Pap tests, HPV testing, and visual inspection are prone to variability, which may result in incorrect diagnosis. The goal of this project is to use deep learning methods to create an integrated system that will automatically detect the type of cervix and classify cervical cancer. A dataset of 915 histopathology images and 4005 colposcopy images was utilised to train and compare several pre-trained algorithms. For the extraction of the area of interest, a lightweight MobileNetv2-YOLOv3 model was used, and for the categorization of cervix types, EfficientNetB0 was employed. The findings revealed that the mean average accuracy for ROI extraction was 99.88%, while the test accuracies for cervix type and cervical cancer classification were 96.84% and 94.5%, respectively.

Vijayanand [16] a novel technique for precisely identifying and categorising Pap smear cell images—which are essential for determining cervical cancer—is presented in this research. A modified deep learning technique based on dual-tree complex wavelet transform (DTCWT) is used in this approach to categorise images into four categories: normal, dysplastic, superficial, and cancer in situ. Three primary modules comprise the method: one for data augmentation, one for DTCWT, and one for ConvNet. For the ConvNet module to achieve high classification accuracy, a large number of cell images are needed. DTCWT is used to convert the enhanced images into multimodal pixels, producing matrix-based sub-band coefficients. ResNet 18 is then used to train and classify these matrices, assigning one of the four classes to the original Pap smear cell image.

Omneya Attallah [17] presented a CAD model that does not need a pre-segmentation procedure by extracting features from many domains. Unlike existing approaches that employ single DL models, the model combines three compact deep learning models to collect high-level spatial deep information. It provides a thorough depiction of the characteristics of cervical cancer by combining statistical and textural descriptors from a variety of domains, such as the spatial and time-frequency domains. The influence of handmade characteristics on diagnostic accuracy and the ramifications of combining combined handcrafted features with DL feature sets from multiple ConvNets are examined in this study. The whole DL features are fused with combined handmade features using principal component analysis, resulting in 100% accuracy with the quatric SVM classifier.

Keshav Kaushik [18] digital imaging and machine learning are being used by clinicians to enhance cervical cancer screening. Although women who test positive for HPV usually have little risk, it is important to identify these women so that they may get treatment right once. Using deep learning methods, researchers have created a model for predicting cancer risk. The model is trained to predict cervical cancer by importing datasets, standardising, and visualising the data. The Cervical Cancer dataset is used to assess the model’s performance and accuracy. This method might assist in identifying women who are at risk of cervical cancer and has important clinical implications.

Madhura Kalbhor [19] the approach for predicting cervical cancer using pap smear images is presented in this research. It uses machine learning methods to create pre-trained deep neural network models, which are then used for feature extraction. For feature extraction, four pre-trained models are optimised: AlexNet, ResNet-18, ResNet-50, and GoogLeNet. After that, the technique employs a variety of machine learning methods. When combined with the AlexNet pre-trained model, basic logistic regression yields the greatest accuracy of 95.14%. This strategy is essential for lessening the arbitrary character of cancer diagnosis. Samia M [20] bilinear pooling is investigated as a way to enhance classification accuracy and expedite processing times in ConvNets. It presents a dyadic feature extraction procedure that creates composite feature representations by multiplying elements-wise and using twin feature extraction modalities. After that, dimensionality is decreased by a random projection process, producing an image descriptor that is clearer and more useful. The RP-BCNN framework obtains an accuracy of 0.9530 for multiclass classification with seven different labels and 0.9983 for dual-label classification situations.

Javier Civit-Masot [21] HPV-related cervical cancer needs molecular-based detection. In spite of immunisation campaigns, incidence rates are greater in places with little access to healthcare. Convolutional neural network classifiers and liquid cytology are employed for mass screening; nevertheless, pathologist verification is necessary for the final diagnosis. Using liquid-cytology images, this research created a bespoke deep-learning classifier to differentiate between four cervical cancer severity groups. With execution speeds under one second, the final classifier obtained over 97% accuracy for four classifications and 100% accuracy for two classes. This study yields better outcomes than earlier studies at a lower computing cost.

Abinaya [22] a cervical cancer categorization method based on deep learning has been developed to tackle this worldwide health concern. The system extracts spatiotemporal characteristics from cervical images using a Vision Transformer module and a 3D convolutional neural network. The design of the system consists of an input layer that receives images, a 3D convolution block that extracts features, and downsampling that removes unnecessary information. Feature maps at different levels of abstraction are extracted using four different Vision Transformer models. Within the 3D feature pyramid network (FPN) module, the feature maps are concatenated, and a 3D average pooling layer minimises dimensions. A kernel extreme learning machine (KELM) is then used to classify the output into one of five groups. The simulation findings demonstrate an outstanding 98.6% accuracy, indicating that it is a potential technique for accurate categorization of cervical cancer. Its potential for use by medical professionals as a diagnostic tool is also clear. Sher Lyn Tan [23] the goal of this project is to create deep learning models that can automatically diagnose cervical cancer without the need for special features or segmentation techniques. Owing to a lack of data, pre-trained ConvNet models were used to apply transfer learning to Pap smear images for a seven-class classification job. Thirteen pre-trained deep ConvNet models were thoroughly evaluated and compared using Google Collaboratory’s Keras package and the Herlev dataset. When it came to accuracy and performance, DenseNet-201 was the best model. The ConvNet models that were pre-trained produced encouraging results in experiments and used less computing time.

Jesse Jeremiah Tanimu [24] more than 85% of cases of cervical cancer occur in underdeveloped nations, making it one of the leading causes of early death. This study proposes a prediction model that makes use of screening and medical record risk trends. The model analyses risk variables related to cervical cancer using a decision tree (DT) classification technique. The model identifies critical characteristics for cervical cancer prediction using least absolute shrinkage and selection operator (LASSO) and recursive feature elimination (RFE) approaches. The SMOTETomek method is used since the dataset has a significant degree of imbalance and missing values. The usefulness of feature selection and class imbalance reduction is shown by comparative analysis. The DT classifier achieves a sensitivity of 100% and an accuracy of 98.72% when paired with certain characteristics. Anurag Tripathi [25] cervical cancer is the fourth most common illness among women worldwide. Recovery rates have been shown by early discovery and early surveillance; nevertheless, morphological alterations make it difficult to differentiate cervical cells in Pap smear testing. For a painless and cost-effective diagnosis, Pap tests are recommended. This research presents ResNet-152 architecture-based deep learning classification techniques for Pap smear images. With a high classification accuracy of 94.89%, the method establishes a standard for further classification methods. Table 1 illustrates the various state-of-the-art model’s outcome comparison along with limitations.

**Table 1.** Model comparison of state-of-the-art methods.

| Author | Year | Dataset | Model | Limitation |
| --- | --- | --- | --- | --- |
| [11] | 2022 | UCI repository | SVM, XGBoost | Computational complexity, potentially limiting their scalability and usability. |
| [12] | 2021 | Maternal and Child Hospital of Hubei Province | ConvNet | Annotated data from multiple hospitals and imaging instruments for training and validation. |
| [13] | 2021 | SIPaKMeD | VGG | System's effectiveness may be constrained by potential limitations inherent to this specific architecture. |
| [14] | 2022 | CRIC | SVM | Its lack of validation on external datasets or in real-world clinical settings. |
| [15] | 2022 | SIPaKMeD | MobileNetv2-YOLOv3 | The reliance on a specific dataset for training and validation, which may introduce biases and limit its generalizability to diverse populations and settings. |
| [16] | 2021 | Herlev | DTCWT, ConvNet | The proposed method is its reliance on a specific dataset for testing, which may not fully represent the variability and complexity of Pap smear cell images encountered in real-world clinical practice. |
| [17] | 2023 | SIPaKMeD | ConvNet, SVM | The proposed CAD model is its reliance on a single type of machine learning algorithm, specifically the quartic SVM, for classification. |
| [18] | 2022 | SIPaKMeD | ConvNet | The proposed cancer risk prediction model is its reliance on deep learning without considering potential biases in the dataset or the interpretability of the model's predictions. |
| [19] | 2023 | SIPaKMeD | Alexnet, Resnet-18, Resnet-50 and GoogLeNet | The proposed methodology is the lack of diversity in the pre-trained models used for feature extraction. While AlexNet, ResNet-18, ResNet-50, and GoogLeNet are well-known deep neural network architectures. |
| [21] | 2023 | Mendeley liquid cytology dataset | ConvNet | The proposed classifier is its reliance on liquid cytology images alone for distinguishing between severity classes of cervical cancer. While liquid cytology is a valuable screening tool, it may not capture all relevant features or abnormalities indicative of cervical cancer. |
| [22] | 2024 | SIPaKMeD | 3DCNN | The proposed DL-based cervical classification system is its reliance on simulated data for evaluating its performance. While simulation results may provide valuable insights into the model's behavior and capabilities. |
| [23] | 2024 | MDE-Lab | DenseNet-201 | The study is the reliance on transfer learning with pre-trained ConvNet models, which may not fully leverage the unique characteristics and intricacies of cervical cancer pathology. |
| [24] | 2022 | ML repository as Risk Factors dataset | DT, LASSO) | The study is that it focuses solely on the application of decision tree (DT) classification algorithm without exploring the potential benefits of other machine learning techniques. |

## 3. Materials and Methods

### 3.1 Materials

The 400 Pap smear images that make up the CRIC dataset have each been painstakingly annotated to identify 11534 cells. These images fall into several categories: possibly non-neoplastic (ASC-US), atypical squamous cells of undetermined significance (ASC-H), squamous cell carcinoma (SCC), low-grade squamous intraepithelial lesion (LSIL), high-grade squamous intraepithelial lesion (HSIL), and atypical squamous cells. Furthermore, this article uses only cropped images (4789 total) that display cervical cells [26]. A preview of the image samples from the CRIC dataset may be seen in Figure 1. The SIPaKMeD dataset, which comprises 4049 images taken and cropped using cutting-edge camera technology, is an invaluable tool for research on cervical cell image categorization. Every image offers a different perspective on the appearance and pathophysiology of cervical cells. A systematic framework for study is provided by the five categories into which the dataset is divided: superficial intermediate, parabasal, metaplastic, koilocytotic, and dyskeratotic. This classification enables more in-depth research on traits and variances among cervical cell types. The variety of cervical cell morphologies shown in the collection highlights the depth of data that may be analysed and understood. Researchers may get important insights into cervical pathology and health via this investigation, which will help with medical diagnosis and treatment plans.

Figure 2 depicts the SIPaKMeD dataset images. The goal of the project was to combine the SIPaKMeD and CRIC datasets to provide a thorough framework for examining cervical cell images. The six categories in the CRIC dataset and the five categories added by the SIPaKMeD dataset were combined to create the merged dataset, which increased the study’s scope to 11 unique categories [27]. This increased the variety of cervical cell types that were being studied and added more morphological variants and clinical presentations to the dataset. The combined dataset yielded 8838 images that illustrate the heterogeneous terrain of cervical cell shape and pathology. These images provide important new perspectives on cervical health and illness. A strict data splitting technique was used, with each subset being randomly chosen for testing, validation, and training in order to guarantee generalizability and dependability. This stratified method reduced bias and improved the analysis’s resilience. For the training, validation, and testing sets, we chose 60%, 25%, and 15%, respectively, randomly from each class of images. In order to facilitate a comprehensive comprehension of the experimental framework, Table 1 offers specific configurations of these data allocations. These configurations provide light on the distribution of images across different subsets and categories. Table 2 represents the two-dataset integration value to evaluate the proposed model. Figure 3 depicts the overall dataset utilization value of each class.

**Figure 1.**
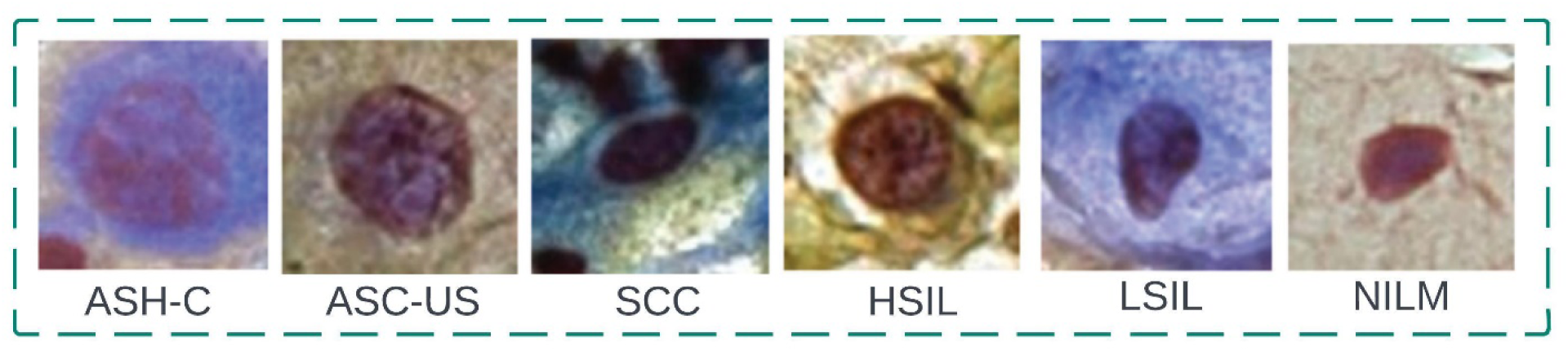
CRIC dataset sample images

**Figure 2.**
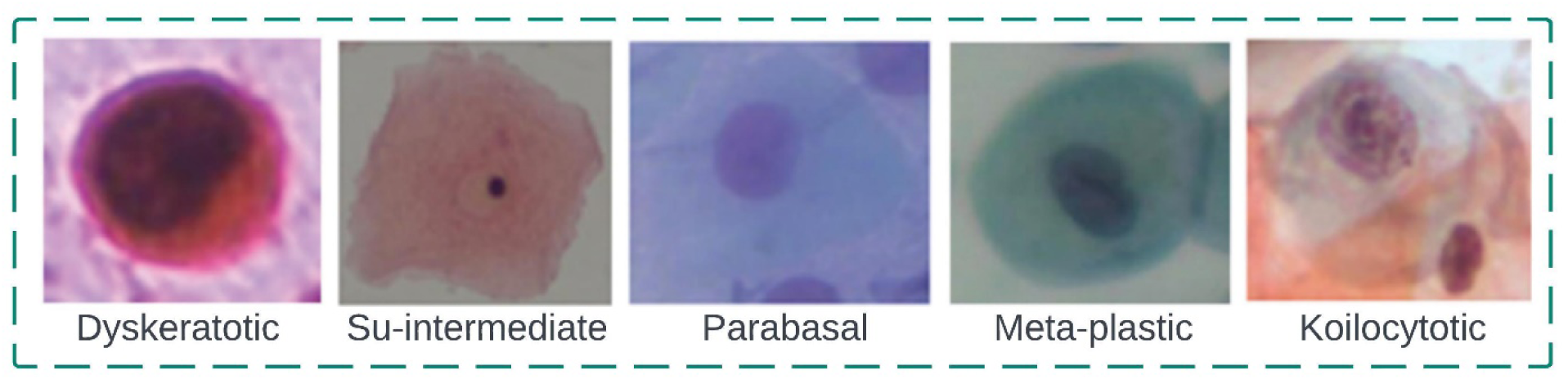
SIPaKMeD dataset sample images

**Table 2.** Integration of two dataset images.

| Class | Training | Validating | Testing | Total |
| --- | --- | --- | --- | --- |
| NILM | 517 | 216 | 129 | 862 |
| LSIL | 491 | 205 | 123 | 818 |
| Koilocytotic | 495 | 206 | 124 | 825 |
| Parabasal | 472 | 197 | 118 | 787 |
| Su-Intermediate | 499 | 208 | 125 | 831 |
| Metaplastic | 476 | 198 | 119 | 793 |
| Dyskeratotic | 488 | 203 | 122 | 813 |
| SCC | 412 | 172 | 103 | 687 |
| ASC-H | 484 | 202 | 121 | 806 |
| HSIL | 524 | 219 | 131 | 874 |
| ASC-US | 445 | 186 | 111 | 742 |
| **Total** | **5303** | **2210** | **1326** | **8838** |

**Figure 3.**
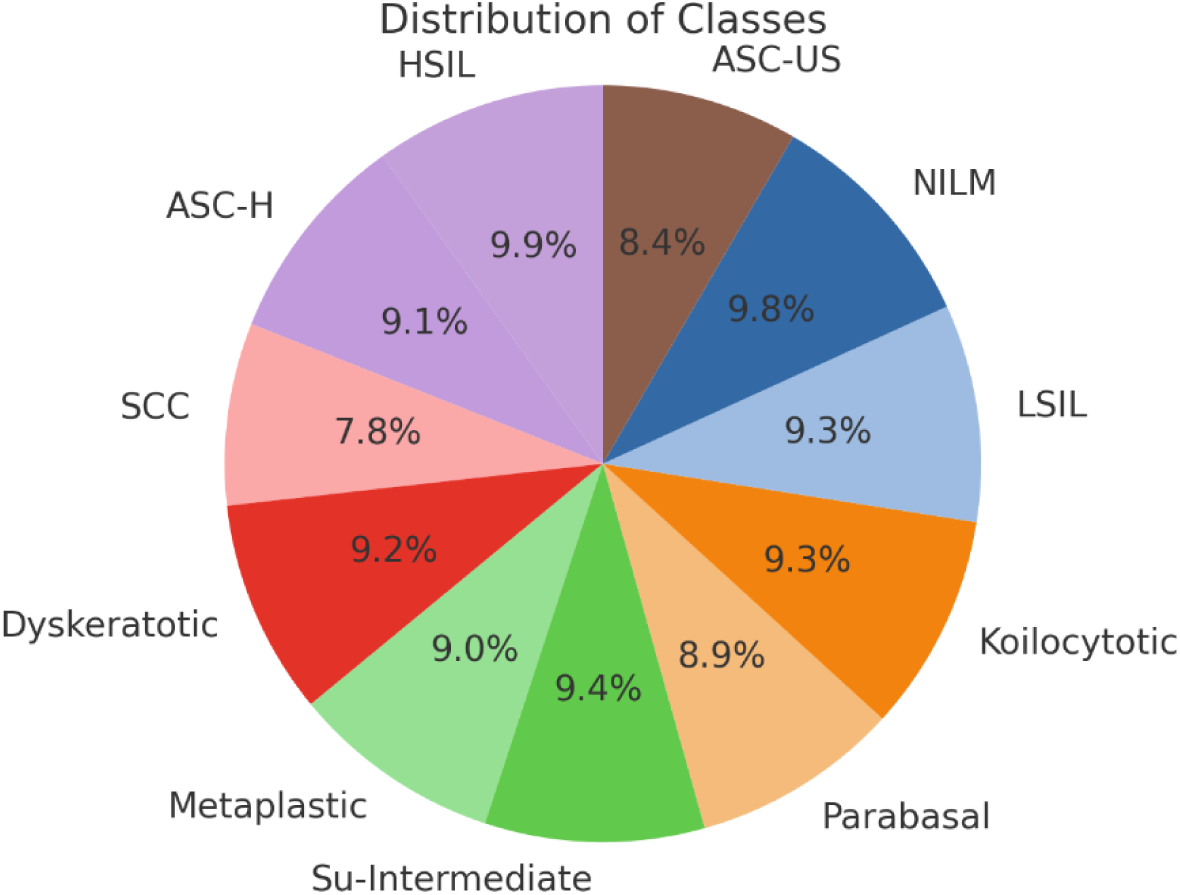
Dataset class distribution overall for 100%

### 3.2 Methods

The purpose of this study is to develop novel approaches in order to enhance the categorization of cervical cancer, which currently occurs in eleven distinct forms. In this study, a complete collection of images was obtained from the CRIC and SIPaKMeD archives. The images were then subjected to a careful preprocessing procedure in order to improve contrast and clarity. It has been hypothesised that the SQ-ResNet152 classification model, which is founded on the concepts of transfer learning, may identify small changes across a number of different forms of cervical cancer. The optimisation of the model’s hyperparameters, which may be accomplished with the help of the DHO optimizer, is absolutely necessary for the success of the model. By adjusting these parameters to perfection, it is possible to maximise the performance and effectiveness of the model, which will eventually lead to advancements in the area of cervical cancer detection and therapy. With the help of this research, the accuracy of cervical cancer detection and therapy will hopefully be improved. Figure 4 depicts the operation flow of the proposed model with transfer learning technique. Preprocessing involves preparing the data for training by doing operations such as resizing images, converting formats, and normalising pixel values to guarantee consistency and interoperability. After that, test images are subjected to analysis using a series of convolutional, pooling, and fully connected layers in a ConvNet model. This results in the production of probabilities that represent the image’s affiliation with different classes. Transfer learning may also be used to improve performance by starting the model using weights that have already been learnt from a ConvNet that has been trained on large amounts of data. Hyperparameter tuning then becomes crucial as changes are made to parameters such as the number of layers, learning rate, and optimizer to maximise model accuracy. This process is then verified against an independent test set to see how well the model generalises to new data.

**Figure 4.**
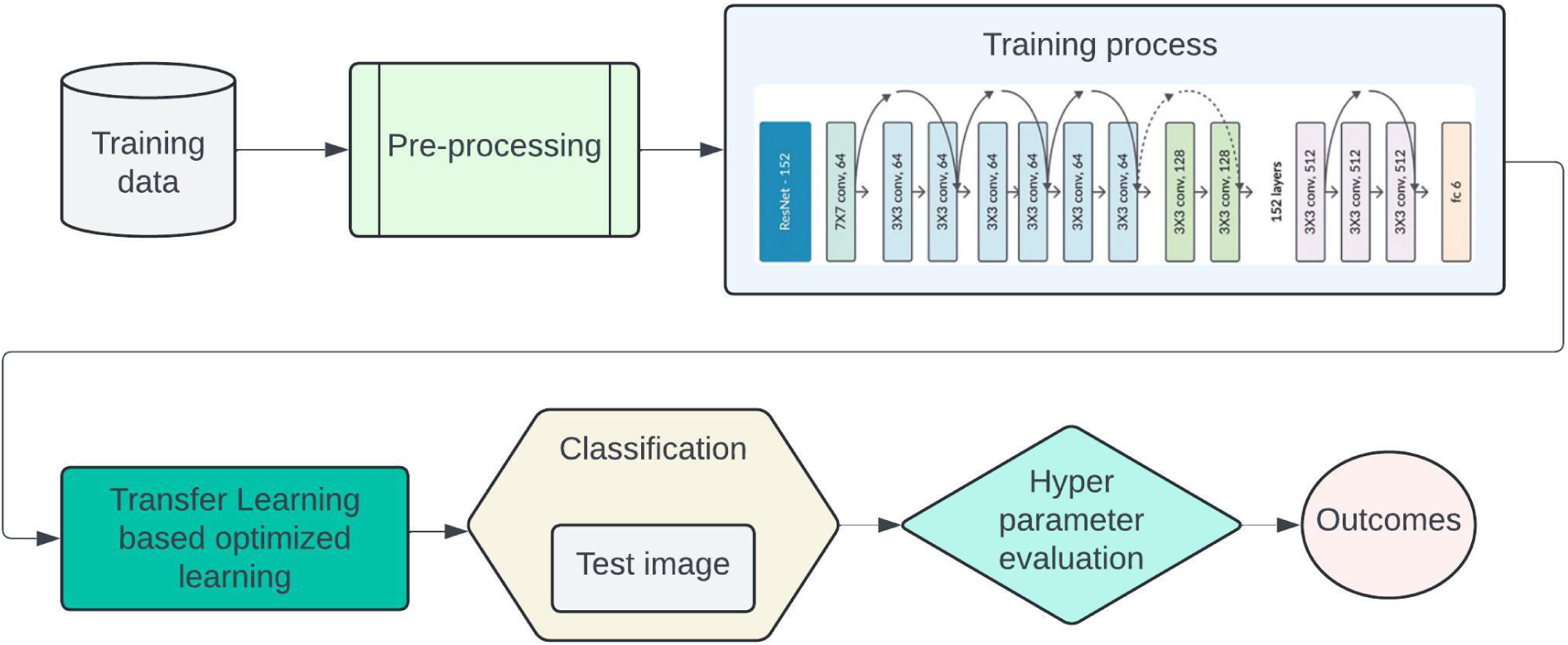
Schematic diagram of proposed work

#### 3.2.1 Pre-processing

In this research, the pre-processing of cervical cancer images is very important as it facilitates the appropriate preparation of the data that will yield the best model. The first operation here involves the change of scale two-dimensional images into equal measurement so that there is uniformity across the dataset. This procedure is important in ensuring that the input specifications of the ConvNet architectures like ResNet152 are adhered to. The JPG or PNG formats are also used on the images creating a homogenous dataset as it was before all the images are then formatted to one extension. Then, pixel intensities in every image are also moved to an appropriate range for instance [0,1] in order to comply with the weights of the model that has already been trained and for the effective training stage with precision. Moreover, the contrast of many images is adjusted using enhancing methods like histogram equalization to make sure that the important features used in diagnosing different types of cervical cancer can easily be seen. In some instances, it was necessary to carry out denoising with Gaussian blur or median filtering in other to endure while maintaining detail. Also, to reduce overfitting, data augmentation incorporating moderate deep learning techniques like random flipping, rotation, and scattered zooming and brightness changing were implemented. All the above pre-processing steps make sure that the images are fully prepared for efficient processing and classification by the SQ-Resnet152 model. Through the use of individualised contrast enhancement processes, the application of mathematical morphology approaches offers a major step forward in the process of improving the quality of Pap smear images. The approaches in question investigate the underlying structural characteristics of the things that are shown in the images [28]. They make use of the complex interaction between the components and the basic mathematical principles in order to identify and make clear the spatial connections that exist. Our methodology provides a full refining of image characteristics by using two different datasets as inputs. This, in turn, enhances the accuracy of future studies. Equation (1) and equation (2) serve as guiding principles within the framework of our morphological operators. They define the erosion and dilation processes with a high degree of accuracy. By performing these procedures, the gray-level image matrix is methodically shaped, and the parameters of the structural elements are meticulously configured in order to maximise contrast and increase the likelihood of proper interpretation.

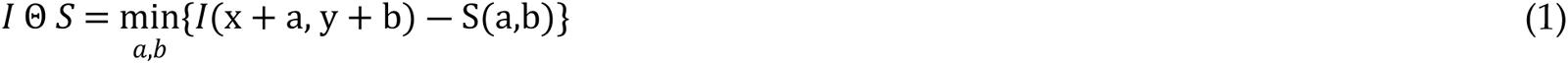

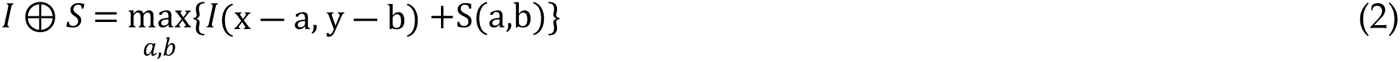

Here, *I*(*x*,*y*) is the intensity of the pixel at position (*x*,*y*) in the image. *S*(*a*,*b*) is the structuring element or kernel, which defines the neighbourhood for the operation. The dimensions of objects diminish as they undergo erosion, and this leads to a progressive reduction in the spatial extent of features within the image. This reduction, which entails the removal of many minute elements and delicate traits, effectively simplifies the overall visual depiction. In contrast, using the dilation operator in reverse causes the objects’ sizes to rise while concurrently reducing the spacing between adjacent features. This increase in object dimensions results in an increased level of prominence and detail added to the image, which also enhances the image’s perceptual clarity. As a consequence, the closure operator is in charge of coordinating the synthesis of the effects produced by the erosion and dilation processes, which leads to a complete refinement of the image structure becoming seen. When these movements are combined, gaps and boundaries are filled in, leading to the final product of a cohesive and well-executed visual composition. Conversely, the opening operator follows the guidelines provided in equation 3, which describes the careful equilibrium that has to be maintained inside the image domain to minimise artefacts and noise while preserving important features.

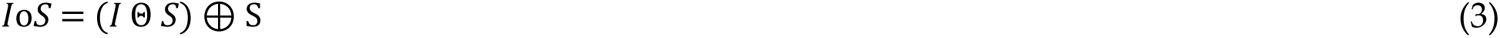

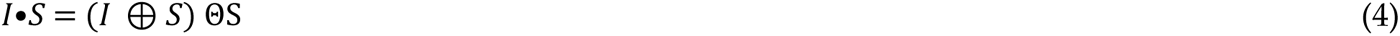

Here, *I* Θ *S* is erosion of the image *I* by the structuring element *S*. Then (*I* Θ *S*) ⊕ S is the dilation of the eroded image by the same structuring element. Then (*I* ⊕ *S*) ΘS is he erosion of the dilated image by the same structuring element. It helps fix and attenuate negative connects that may exist between artefacts and sparse information within the image; it is not sufficient for the opening operator to only fill in the blanks. This is one of the opening operator’s most important tasks. This operator selects and removes undesirable elements while leaving behind important details to provide a polished and coherent visual depiction. This is achieved by the preservation of key elements. Interestingly, disk-shaped structural elements (SEs) are the preferred choice above other types of masks in the field of medical imaging. The fact that disk-shaped SEs are consistent with the anatomical characteristics often seen in medical images may contribute to their widespread adoption. It is feasible to do more intuitive and accurate image analysis because to this compatibility. Even though SEs are becoming more and more prevalent, choosing a size and style is still mostly up to the individual. The specific requirements of the imaging task at hand as well as empirical elements have a role in this selection. This adaptability highlights how morphological processes may be adjusted to a broad range of imaging situations, leading to the creation of highly efficient and uniquely personalised image processing systems.

#### 3.2.2 Feature Extraction – Transfer Learning

Feature extraction is one of the most important steps contributing to the success of the classification problem, especially with transfer learning techniques. It is worth noting that the SQ-ResNet152 model pre-trained on a large dataset such as ImageNet is already able to perform image feature extraction concerning elements like edges, shapes, and textures. Therefore, rather than building the necessary architecture and features from the philtre, we benefit from the available layers and emphasize building features that are more complex than those that are usually relied on capturing whilst training models on Pap smear images. This process begins with the network being able to detect coarser and crude information from the features obtained after the network has been pre-trained and proceeds to modifying the various layers to get deeper level features of the domain. In the case of Pap smear images, consideration is given not only to main textural and statistical characteristics but also to quite sophisticated ones such as wavelet and morphological features essential for cervical cancer detection. The model changes the pattern of feature extraction that includes the previously trained elements to include aspects that are unique to this classification problem, thereby obviating the need for retraining and a lot of data. The concept of transfer learning is rather important in our model since it allows the SQ-ResNet152 to make use of general datasets as a knowledge base and focus on the peculiarities of the task of interest: Classification of Pap smear images. Instead of training the model from scratch which is impossible due to data and computation resource limits, we initialize the model with weights obtained from another network that has been trained with the ImageNet dataset. Such weights are able to retain low level information like images of shapes, edges and textures that are common in most images. While training, instead of considering the model parameters as randomly generated during the commencement, they all begin from these pre-trained weights. The model then goes through a process whereby certain layers are trained more on cervical images obtained from special datasets (the CRIC and SIPaKMeD datasets) allowing it to learn to recognize cervical images. This method enhances the model’s capacity to identify cervical cancer patterns owing to the targeted adaptation of the learned features from the pre-training phase towards the image’s cuts pertaining to the classification task.

Within this framework, transfer learning further reduces the training time of the model by beginning from a well-built model. The SQ-ResNet152 model is already trained and possesses pre-set weights to help detect certain generic features in an image. Because of the fine-tuning conducted on our datasets, the optimization is aimed at the already-existing weights and is more concerned with the features in Pap smear images. This saves a lot of data and time which otherwise could have been used to train the model for top performance. Rather than recommending more exhaustive tuning of hyperparameters from a blank slate, the approach requires this tuning of parameters with respect to the pre-trained model but rather for the task. The pre-trained model understands the general structure of the images which leads to faster convergence and better performance.

##### 3.2.2.1 ResidualNetwork152

The use of residual adding units stands out as a particularly significant method for dealing with the problem of disappearing gradients, which is one of the fundamental problems in the area of deep neural networks [29]. This innovative approach has changed the game by effectively overcoming the obstacles that had previously prevented deep learning from progressing. Specifically, ResNet has been the impetus for a paradigm shift in image identification, ushering in a new era that will stand out for its creative methodology. However, despite these amazing developments, it is still impossible to overstate the importance of depth in determining how successful feature learning is. We may unlock the potential to delve deeper into the intricacies of image representation by stacking layers within the neural network design. This enhances the complexity and depth of the attributes that are retrieved. In our study, we use the power of a state-of-the-art ResNet variant called ResNet152, which has an astounding 152 layers. This robust design serves as the foundation of our method and allows us to extract cervical imaging data with remarkable precision and accuracy. In addition to increasing feature extraction accuracy, ResNet152’s greater depth enables us to identify minute variations throughout the Pap smear image collection. Acknowledging these nuances is a noteworthy achievement.

**Figure 5.**
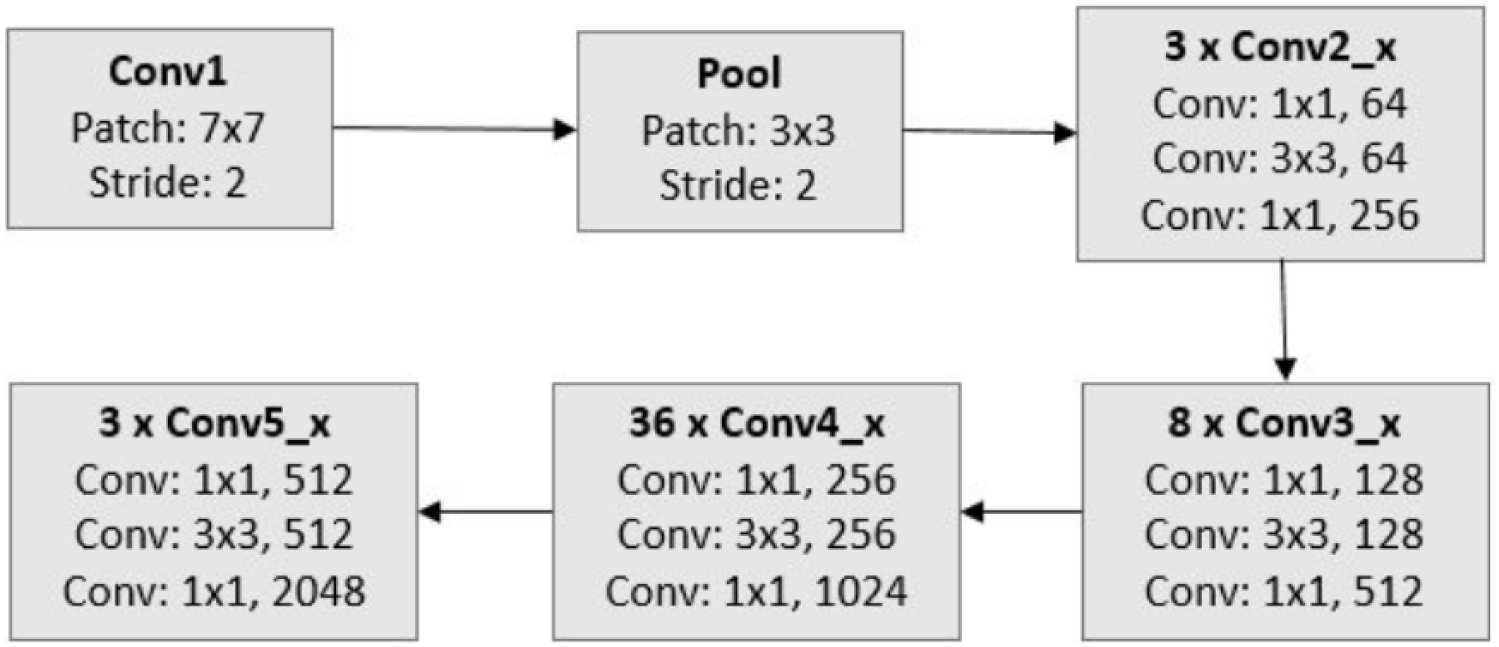
ResNet152 building block

Specifically, the model does very well in capturing the small-scale and fine-grained features seen in cervical images. It thereby establishes the foundation for enhanced categorization performance. Our goal is to advance the detection of cervical cancer by using a combination of accurate feature extraction techniques and advanced neural network architecture. More precisely, we want to provide an overview of deep learning’s transformative potential for applications in medical imaging. Figure 5 depict the basic building blocks of ResNet152. The residual operating is represented as,

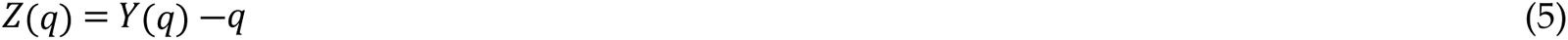

During the training phase, the parameter function, which is denoted by the letter *Y*, is learnt directly. The input is *z*, and the residual function is represented by the symbol *Z*(*q*). When it is possible for ‘simple’ layers to directly accept a putative identity mapping function *Y*(*q*) = *q*, training deep neural networks becomes a process that is difficult to do. On the other hand, the residual network provides a solution that is more feasible, optimising and converging at a faster rate. This is accomplished by translating *Y*(*q*) into a residual function, which is written as *Z*(*q*) + *q*. Taking this method not only simplifies the optimisation process but also makes it easier to achieve convergence more quickly. Through the process of gradually increasing the depth of the network, we are able to unleash the potential for increased feature extraction capabilities. This enables the network to catch patterns within the data that are more subtle and complicated.

##### 3.2.2.2 Squeeze-and-Excitation

The provision of Squeeze-and-Excitation (SQ) module modifies the use of the provision of medical imaging applications whereby the detection of cervical cancer enhancement needs a more accurate feature representation. In this context, SQ module aids the neural networks in remembering the key features in cervical cancer screening images such as the critical patterns of a tumor such as its shape, size, and texture. Figure 6 depicts the basic SQ model representation.

- Squeeze: the high dimensional images in this scenario, are medical images which may include finer details like cell structures, tissue malformations and tumour margins and such details are processed in the network. In this phase, the SQ module handles the problem of how further processing should reduce the spatial complexity of these images by including important channel-wise attention throughout the image. This is critical as it allows the network to understand the broad contextual information with respect to the regions with the potential of containing cancer, which is central to detecting the subtle regions indicating cervical cancer.
- Excitation: After global information has been captured, the SQ module learns which channels (features) are most relevant for detecting cancer. For example, it may give more importance to channels that represent certain textures or tissue irregularities, which are common signs of malignancy. In attending to the channel dependencies, the network is able to highlight features that are diagnostic thereby enhancing its performance in the discrimination of cancerous and non-cancerous tissues.
- Scale: The previously adjusted weights of each channel, in regards to the importance of the discriminative feature maps, is used over the raw feature maps. This process of undoing the changes made ensures that the network gives more stress to the most salient features which are the nature and the arrangement of the atypical cell growths which would increase the accuracy level in cervical cancer screening.

However, SQ modules can effectively increase the ability of neural networks in cervical cancer detection systems working with IB subconsciously, detecting time critical pictorial structures assisting ED better, and diagnosing all the disease-related symptoms accurately. Such technology may help doctors provide more accurate image-based predictions for diseases and improve patients’ prognosis.

**Figure 6.**
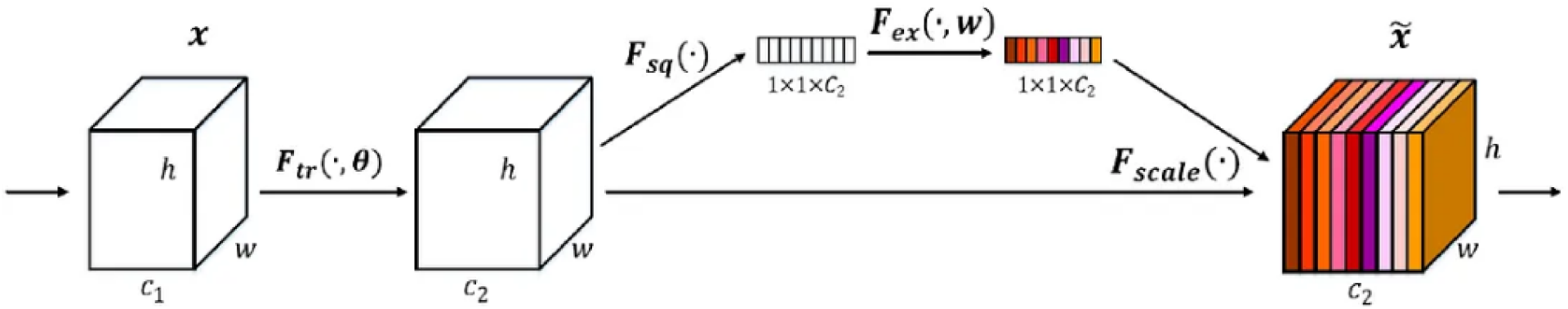
Schematic diagram of SQ model

##### 3.2.2.3 SQ-ResidualNetwork152

The Deer Hunting Optimization Algorithm (DHOA) is an advanced metaheuristic inspired by the cooperative hunting behavior of deer, offering a robust framework for hyperparameter optimization by effectively balancing exploration and exploitation of the search space. In DHOA, hyperparameters are encoded as vectors representing continuous or discrete values, with each candidate solution corresponding to a deer’s position in the search space [30]. The algorithm evaluates these candidate solutions using a fitness function, defined in this study as the negative cross-entropy loss, which balances model accuracy and confidence while incorporating regularization terms to penalize overly complex models for better generalization. The optimization process begins with the random initialization of a population of deer across the search space, with each assigned a fitness value based on the fitness function.

**Figure 7.**
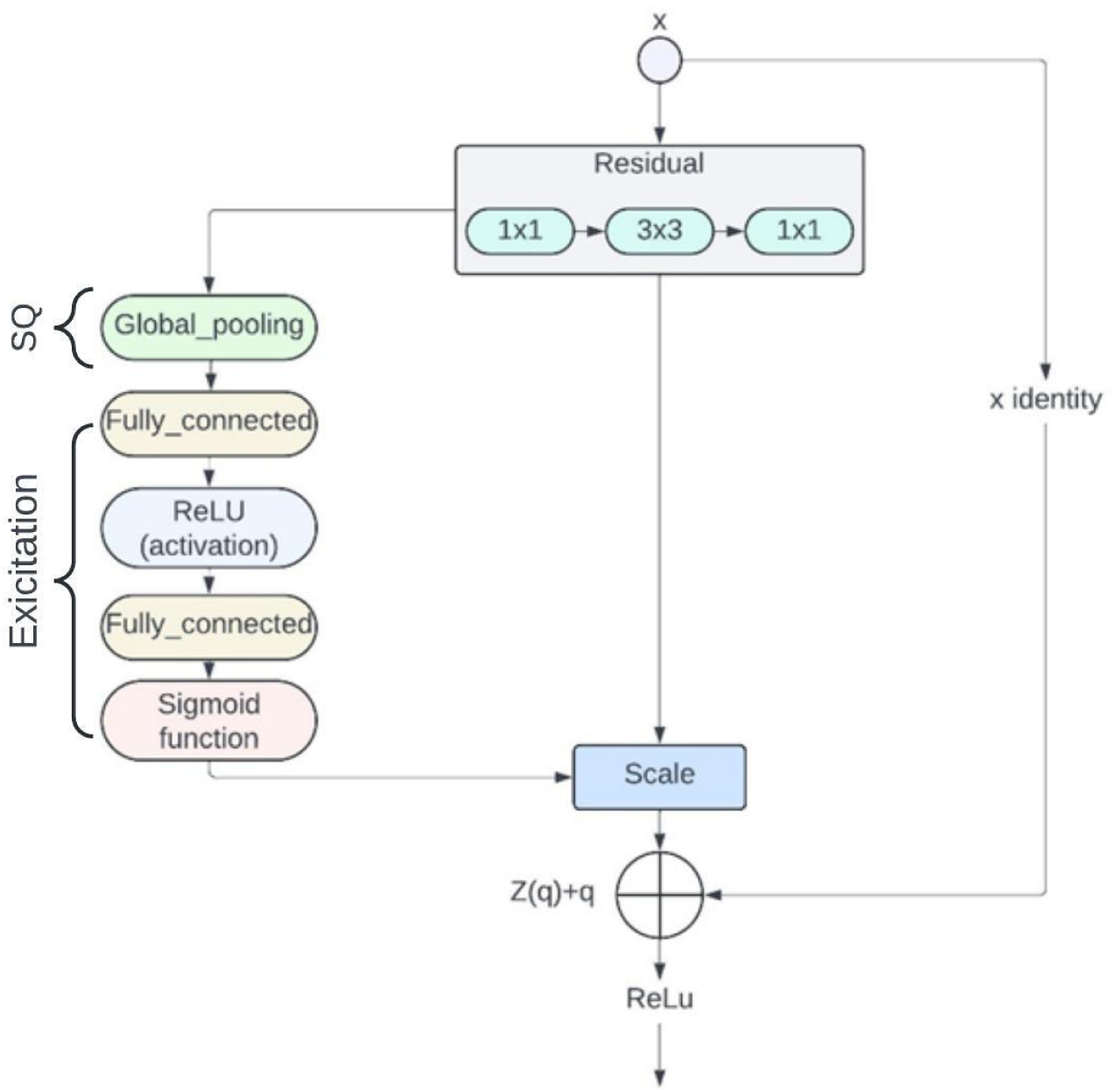
SQ-ResNet152 model structure schematic diagram

The algorithm employs two key strategies to refine solutions: “chasing prey”, which exploits local regions based on nearby solutions’ fitness, and “avoiding predators”, which introduces random perturbations to diversify the population and escape suboptimal solutions. These behaviours, inspired by real-world hunting strategies, ensure both thorough exploration and focused exploitation of the search space. The algorithm iterates through these phases until convergence criteria, such as a maximum number of iterations or a predefined performance threshold, are met. By integrating DHOA into the SQ-ResNet152 architecture, the model achieves optimized hyperparameters, significantly improving its sensitivity to feature variations in cervical cancer image analysis. Figure 7 provides a clear schematic representation of the DHOA process, illustrating the encoding of hyperparameters, the interaction between exploration and exploitation phases, and the evaluation of the fitness function, thereby showcasing the algorithm’s effectiveness in enhancing classification tasks.

**Table 3.** ResNet152 finetuned parameter settings.

| Layer name | Output size | 152-layer |
| --- | --- | --- |
| conv_1 | 112×112 | 7×7, 64, stride 2 |
| conv_2x | 56×56 | 3×3 max_pool, stride 2 |
|  |  | [1×1, 64 @ 3×3, 64 @ 1×1, 256] × 3 |
| conv_3x | 28×28 | [1×1, 128 @ 3×3, 128 @ 1×1, 512] × 8 |
| conv_4x | 14×14 | [1×1, 256 @ 3×3, 256 @ 1×1, 1024] × 36 |
| conv_5x | 7×7 | [1×1, 512 @ 3×3, 512 @ 1×1, 2048] × 3 |
|  | 1×1 | avg_pool, 1000-d fc, softmax |
| FLOPs |  | 11.3×10<sup>9</sup> |

As stated in Table 3, the Image is applied to several layers of neural network that modify the dimensions of the image as follows: The input image has an initial dimension of 112×112 pixels after been subjected to the first convolutional layer convection This procedure utilizes a kernel of 7×7 with a stop of 2, reducing the image size. Moving from layer-to-layer image and its size is getting smaller. After the second layer (conv_2x), max pooling operation reduces the output size to 56×56 with the stride of 2. In addition, there is further down sampling in conv_3x where the size is 28×28. Conv_4x reduces the size to 14×14 and finally conv_5x reduces the image to sizes of 7×7. After the last convolutional layer of processing, a global average pooling reduces the feature map to one instead of 1 not only 1 but with the help of the fully connected layer classification is done. This kind of stepwise shrinking of the image is useful in making the network extract more advanced patterns from the image and overall decrease images spatial redundancy.

##### 3.2.2.4 Loss Function

In cases where data is imbalanced, classifiers often struggle to achieve high classification results. This happens when the number of samples in certain categories of the dataset is much smaller than in other categories. This mismatch may cause the learning process to become skewed, which might provide worse than desirable outcomes, particularly for the underrepresented groups. However, re-weighting the loss function seems to be a powerful technique that may be used to lessen this challenge and increase the diagnosis accuracy of cervical cancer. This method gives the minority classes more weights and the majority classes lesser weights, thus balancing the effects of each group throughout training. By giving the minority groups more weights, this is achieved. The classifier’s ability to accurately detect cervical cancer across all categories eventually improves as a result of its increased ability to dedicate time and resources to the process of learning the complexity of the underrepresented classifications.

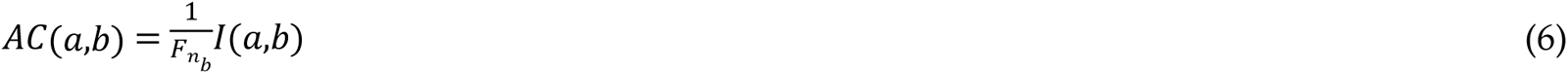

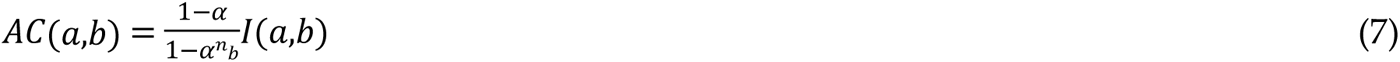

When *S* is the entire sample size of the ground-truth class, represented by *n*_*b*_, and *x* is a specific sample, the loss function defines the probability distribution of the model, represented by *A* = [*a*1, *a*2,…, *Ac*], where each ai ∈ [0,1]. The class-balanced term is adjusted by the hyperparameter ’α’. Equation (9) expresses the class-balanced focal loss.

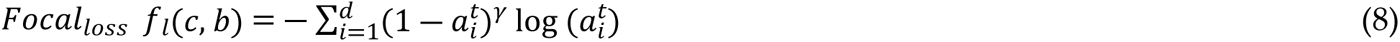

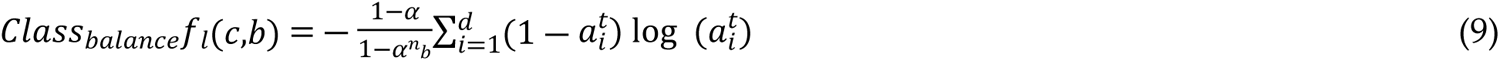

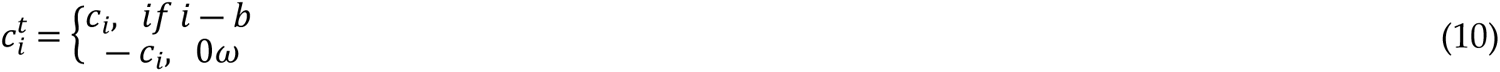

The model’s output is *c* = [*c*_1_,*c*_2_,*c*_3_,….*c*_*d*_]^*T*^ here the overall number of classes is indicated by *C*.

##### 3.2.2.5 DHOA

The Deer Hunting Optimization Algorithm (DHOA) is a unique approach based on deer hunting that is a type of optimization algorithm. While traditionally grounded in understanding the interactions between the hunter and the deer’s natural defense, its computational adaptation focuses on finding the most optimal solution in complex search spaces. In this algorithm, the distinct abilities of deer, such as detecting subtle odors, faint motions, and high-frequency sounds, are modelled as strategies that enhance exploration and exploitation capabilities in the search space. For hyperparameter tuning, DHOA proves particularly effective due to its dynamic balance between exploration (searching new areas in the search space) and exploitation (refining solutions in promising regions). Hyperparameter encoding in DHOA involves representing each hyperparameter as a vector, where each dimension corresponds to a specific parameter (e.g., learning rate, weight initialization, activation function). Each candidate solution is considered a “deer” in the population, with its fitness value determined by evaluating the detection rate on the training dataset. This evaluation uses a carefully designed fitness function, such as negative cross-entropy loss, which ensures the model not only achieves high accuracy but also avoids overfitting. The DHOA optimization process starts with the initialization of a population of “deer” randomly distributed across the search space. During the exploration phase, the algorithm mimics “chasing prey” by focusing on promising regions of the search space based on fitness evaluations. Meanwhile, the exploitation phase introduces random perturbations to simulate “avoiding predators”, enabling the model to escape local optima and maintain diversity. This iterative process continues until the convergence criteria, such as a maximum number of iterations or an acceptable performance threshold, are met.

**Figure 8.**
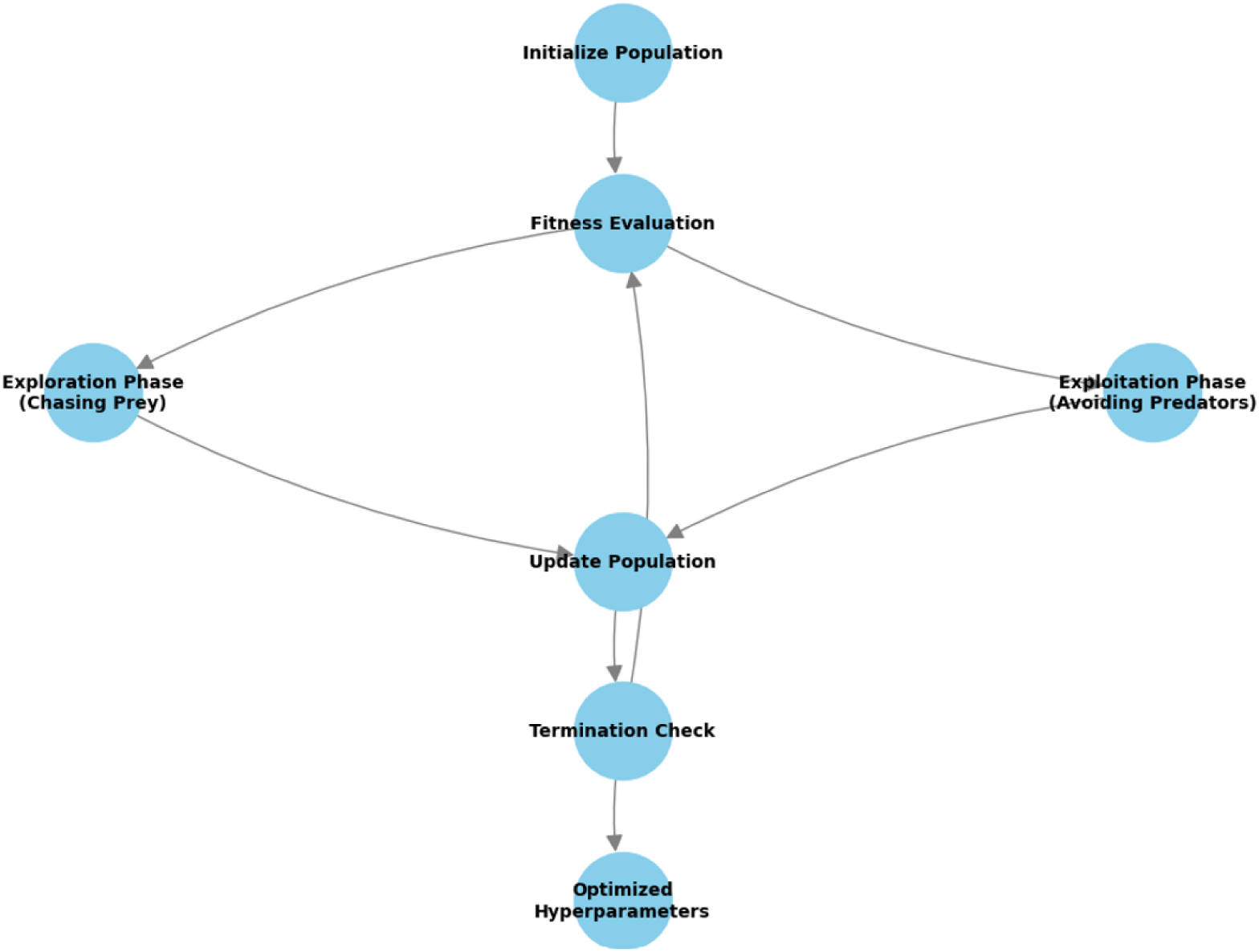
Schematic representation of DHOA

Figure 9 depicts the step-by-step operational flow of DHOA. This, in turn, has an effect on the amount of time required for training and inference, which typically increases as the number of neurons in the network increases. One of the most important factors that determines the capacity of the multilayer model is the number of neurons that are dispersed across its layers. The incorporation of extra layers is beneficial in the context of automated feature engineering since it makes it possible to represent data in a more complete manner and fosters increased learning capabilities. In many instances, the choice to augment the network with extra layers is correlated with better accuracy, especially when presented with complex data representations. The decision about whether or not to augment the network with additional layers is dependent upon the complexity of the dataset. The standard DHOA, on the other hand, is characterised by four separate mathematical procedures. Initialization of the Population is the first stage, and it is at this phase that the original population of hunters is created. There is a population of hunters, which is marked by the letter *H*, and there is also an average number of hunters, which is denoted by the letter *S* shown in equation (11).

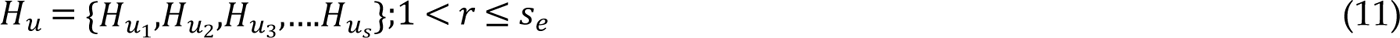

This phase represents the beginning of the process of identifying the best placement for hunting deer. Stage 2 is referred to as Wind Initialization and Position Angle. The wind angle, denoted by φ, and the position angle, denoted by θ, are both determined according to the given information. In addition, a random number that falls within the range of [0, 1] is created, which is indicated by the letter *i*. The current iteration is maintained by the variable *a*, shown in equations (12) and (13).

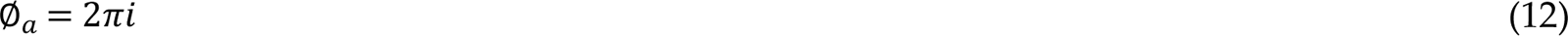

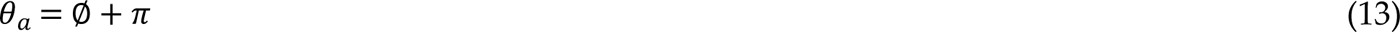

Position Propagation is the third phase, and it is during this phase that the fitness function plays a significant part in determining the most advantageous hunting space, which is the position that is closest to the ideal solution. There are two primary roles that are established, and they are the successor (*H_successor*) and the leader (*H_lead*).

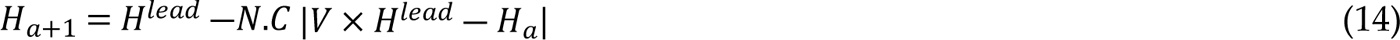

Equation (14) represents the current location of the hunters as *H*_*a*_, and the upgraded position of the predators is described as *H*_*a*+1_ in the subsequent operation. On the other hand, the random number c, which varies between 0 and 2 to account for the wind angle, is shown. This utilises the random integer rd, which spans from 0 to 1. The parameter “*a_max_*” determines the maximum number of iterations, while the coefficient vectors (N) are computed below the equation.

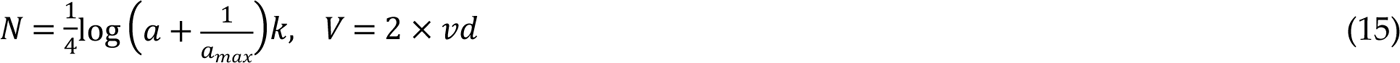

Utilising position angle as a foundation for propagation: In the process of enhancing the hunter’s location, this is achieved by expanding the search area via the consideration of the position angle. The location angle is chosen to enhance the efficacy of deer hunting. Furthermore, the parameter “*deer_a_*” is calculated by subtracting the wind angle from the visualisation angle in order to enhance the accuracy of the position angle, as shown in equation (16). Equation (17) calculates the prey’s visual field angle.

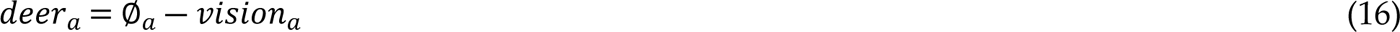

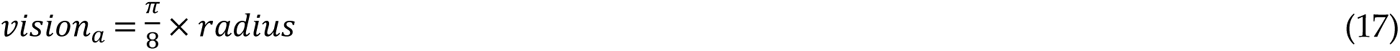

The procedure of propagating the successor position is as follows: The encircling process is used to modify the vector *V* during the exploration phase. The magnitude of the vector *V* is deemed to be less than one when assessing the initial point of the random search. Equation (18) demonstrates how the hunter’s position is updated dependent on the location of the successor. The selection of the search agent is based on a probabilistic approach when the V-value vector is below 1. Otherwise, the best solution is developed to enhance the locations of the search agents.

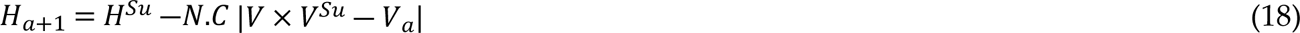

During the propagation process, the leader’s position is used throughout. Once the best roles have been determined, every member of the community makes a concerted effort to obtain them, which kicks off the process of upgrading job descriptions. In this step, the hunter’s encircling formula is guided by the empirical formula from the previous phase. Finalisation is the fourth and last step of the DHOA algorithm. During this phase, the technique for updating the position is repeated and repeated until the optimal hunting site is found.

**Figure 9.**
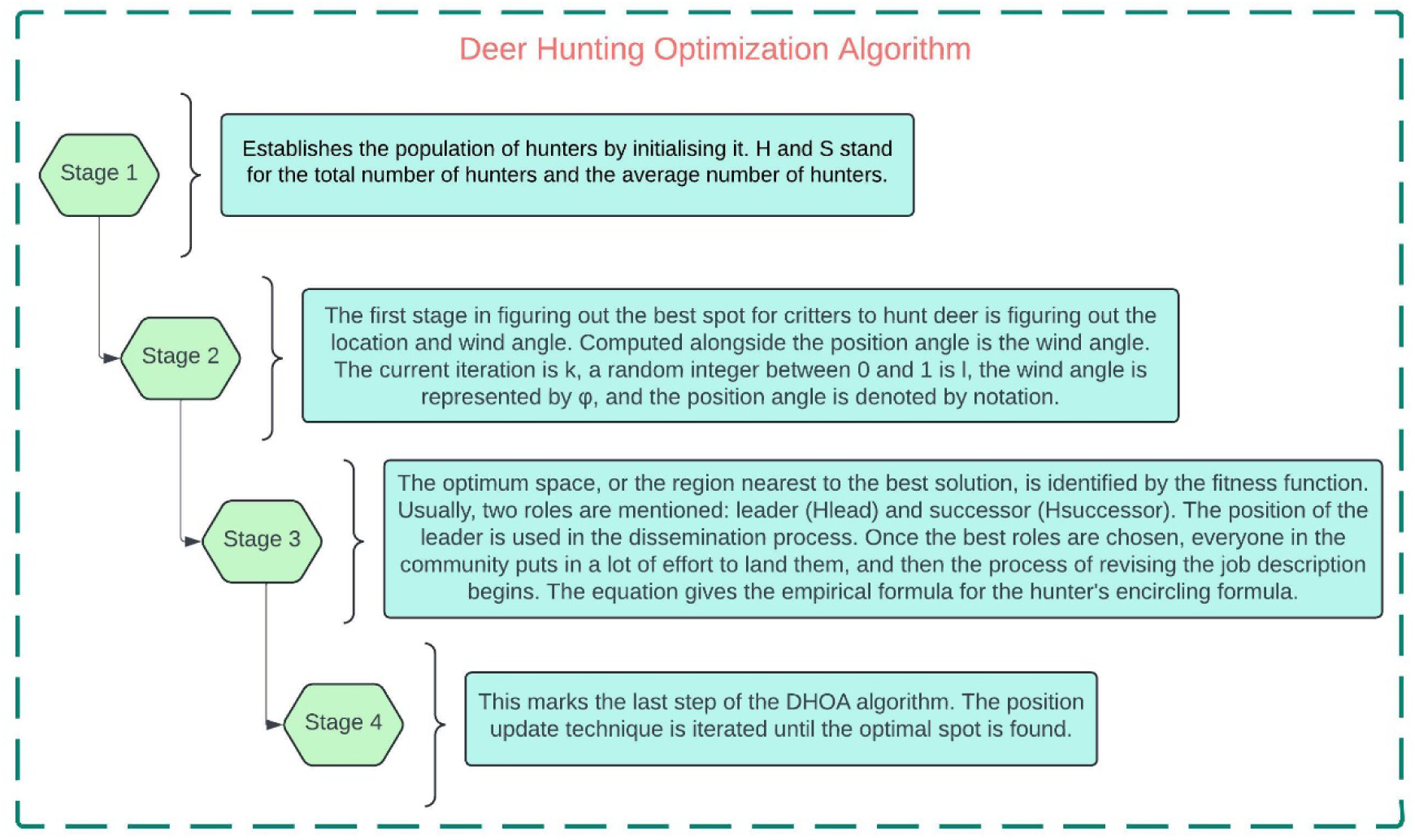
DHOA algorithm overview

## 4. Results and Discussion

This section provides a thorough examination of a suggested method for handling data processing jobs using Python. The study assesses the approach’s efficacy and contrasts it with current cutting-edge techniques. The experimental assessment is conducted using Python, which ensures repeatability and streamlines preparation. The effectiveness of the approach is carefully examined in a range of settings and circumstances. It is essential to standardise picture resolution and size to provide consistency and interoperability across datasets and models. By doing this, biases and inconsistencies are reduced, improving the validity and reliability of the findings. The suggested approach extracts feature and detects patterns using a deep learning architecture called SQ-ResNet152, which is based on transfer learning. The system shows amazing skill in reliably recognising diverse cervical cancer situations and is optimised for cervical cancer classification. To do this, the SQ-ResNet152 method makes use of transfer learning and optimisation approaches, indicating its potential use in real-world clinical scenarios. The comparison study’s findings provide insightful information for further investigations and advancements. The suggested approach shows promise for properly diagnosing and treating cervical cancer situations. It is built on transfer learning, a state-of-the-art deep learning architecture. This method offers promise for practical clinical applications in addition to being successful.

### 4.1 Experimental Configuration

The study experiment was carried out on a local machine that was running Windows 11 and had a capacity of 32 gigabytes of random-access memory (RAM). For the purpose of implementation, the version 1.8.0 of PyTorch was used, which made use of the capabilities offered by the Python programming language. The hyperparameter setup of the Adam optimizer was used during the training of the suggested network model. The specifics of this configuration are shown in Table 4. Key parameters and their corresponding values used in a computational experiment or machine learning model training are shown in Table 4. A momentum setting of 0.8 indicates a rather strong momentum setting. “Momentum” is defined as the percentage of the previous update kept in the current update step. In iterative algorithms, “breakdown”, denoted as 10-4, usually denotes the tolerance level for convergence; in this instance, it denotes a tolerance of 10 to the power of -4. A tiny value of 0.0003 indicates a conservative adjustment strategy. “Rate of learning” indicates the step size or the amount by which model parameters are changed during training rounds. The term “size of batch” describes how many samples are handled during each training iteration; a batch size of 16 is considered to be very minimal. Lastly, “No of epoch” indicates the total number of epochs or training iterations; a value of 100 indicates that a significant number of iterations is required to fully train the model. Together, these parameter values affect the training process’s convergence and optimisation behaviour, which in turn affects the trained model’s efficacy and performance.

**Table 4.** Optimizing hyperparameters for network training.

| Parameter | Value |
| --- | --- |
| Momentum | 0.8 |
| Breakdown | 10<sup>-4</sup> |
| Rate of learning | 0.0003 |
| Size of batch | 16 |
| No of epoch | 100 |

### 4.2 Performance Metric Evaluation

We do calculations based on precision, recall, F1-Score, and accuracy measures in order to assess the performance of the model that has been presented. The number of samples that were properly anticipated to be positive is denoted by the True Positive (T.positive) measure, while the True Negative (T.negative) measure corresponds to the samples that were correctly projected to be negative. False Positives (F.positive) are samples that are negative but are improperly labelled as positive, while False Negatives (F.negative) are samples that are positive but are mistakenly classified as negative. Precision is the ratio of true positives to all samples that were properly predicted, while recall is the ratio of successfully predicted positive samples to all samples that were really positive. A well-rounded evaluation of the performance of the model is provided by the F1-Score, which is a harmonic mean of accuracy (Acc) and recall. It is the percentage of accurately anticipated occurrences to the total number of instances that is referred to as accuracy. These performance measures, when taken as a whole, provide insights into the efficiency and dependability of the predictions made by the model.

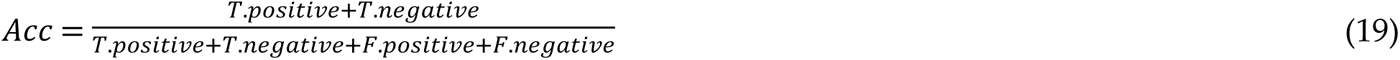

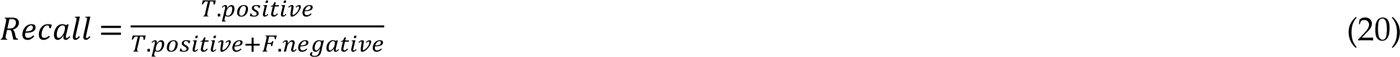

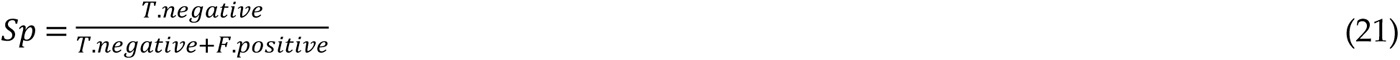

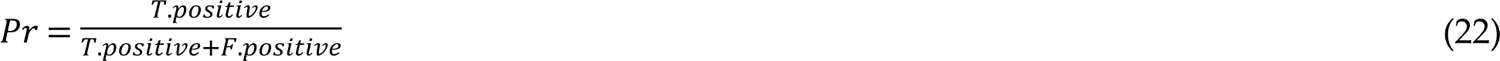

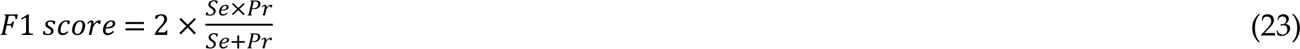

### 4.3 Experimental Result

As shown in Figure 10, the performance of the proposed model is examined throughout the testing and validation stages. This evaluation is carried out using a comprehensive set of metrics that includes accuracy, specificity, F1-Score, recall, and precision. It is clear from the findings that the suggested model is capable of achieving remarkable performance metrics on both the validation and testing sets. More specifically, the model obtains a precision of 98.85%, a recall of 98.63%, an F1-score of 98.84%, an accuracy of 99.69%, and a specificity of 98.76% when applied to the validation set. On the other hand, the model obtains a precision of 98.98%, a recall of 98.15%, an F1-score of 99.06%, a specificity of 98.97%, and an accuracy of 99.79% when it is presented with the testing set. The outstanding generalizability of the model is shown by the fact that there was hardly any discernible change in performance between the validation set and the testing set. In addition, the fact that the suggested model performs better than other techniques that are already in use demonstrates both its effectiveness and its potential to provide exceptional outcomes. Table 5 represents the metric comparison outcome of both validation and testing of proposed model.

**Table 5.** Performance metric comparison of proposed models’ validation and testing outcomes.

| Parameter metrics | Validation (%) | Testing (%) |
| --- | --- | --- |
| Specificity | 98.76 | 98.97 |
| Accuracy | 99.87 | 99.74 |
| Precision | 98.85 | 98.98 |
| Recall | 98.63 | 98.15 |
| F1 score | 98.84 | 99.06 |

**Figure 10.**
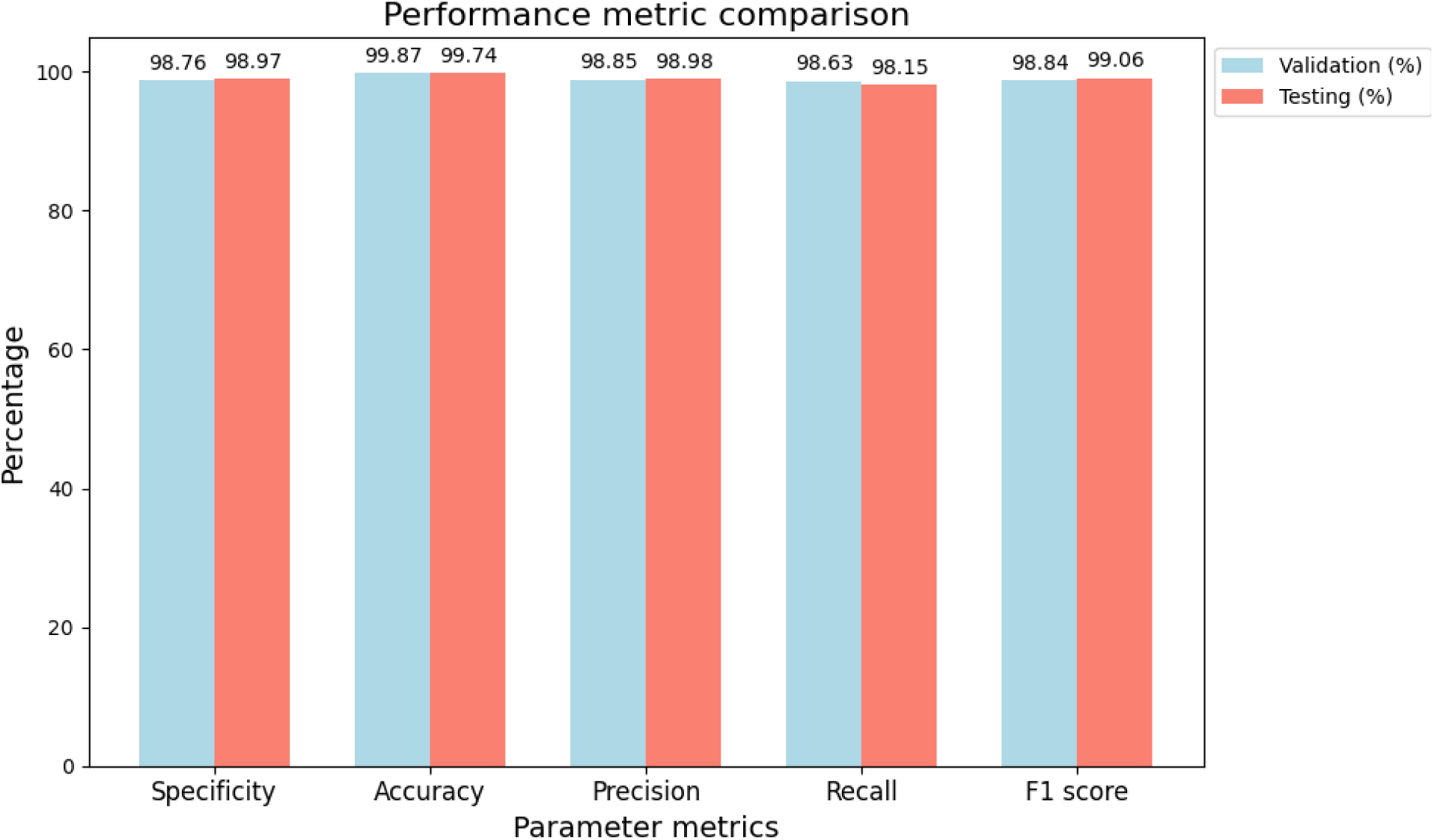
Both validation and testing metric outcome comparison of proposed model

The confusion matrices that are produced by the proposed method for both the validation and testing stages are shown in Figure 12. The confusion matrix for validation provides evidence of accurate categorization results across all categories. For example, fifteen of the images in the second category have been mistakenly categorised, nine of them have been put in the fifth category, and six of them have been placed in the sixth category. In addition to this, twenty-four images that belong to the fifth category have been incorrectly placed to the second category. In a similar manner, the confusion matrix of the test reveals comparable results, with nine images from the second category and twenty-one images from the fifth category being incorrectly classified as belonging to the sixth category. Additionally, eleven images in the fifth category are wrongly classed as belonging to the sixth group, while twenty-two images are incorrectly labelled as belonging to the second category. Both of these errors are quite concerning. In addition, eleven pictures that belong to the eighth category have been incorrectly placed to the seventh category with no explanation.

**Table 6.** Accurately classifying the image based on the category wise.

| Category | Validation | Testing |
| --- | --- | --- |
| 2nd to 5th | 24 | 22 |
| 2nd to 6th | 8 | 6 |
| 2nd to 2nd | 118 | 114 |
| 2nd to 5th | 24 | 22 |
| 5th to 6th | 7 | 5 |
| 5th to 2nd | 15 | 21 |
| 6th to 2nd | 9 | 9 |
| 6th to 5th | 6 | 11 |
| 8th to 7th | 2 | 11 |

In the validation phase confusion matrix, each row represents the true label (or class), while each column corresponds to the predicted label. The matrix shows the counts of instances where a certain true label was predicted as another label. For instance, the first row indicates that there were 151 instances of true label 1, out of which 151 were correctly predicted as label 1, 5 were predicted as label 2, and so forth. Similarly, for the second row, there were 118 instances of true label 2, out of which 118 were correctly predicted as label 2, 24 were predicted as label 5, 8 were predicted as label 6, and so on. This pattern continues for all true labels, providing insight into the model’s performance in the validation phase. Figure 11 (a) depicts the confusion matrix for validation test phase. Figure 11 (b) represents the confusion matrix for testing phase.

**Figure 11.**
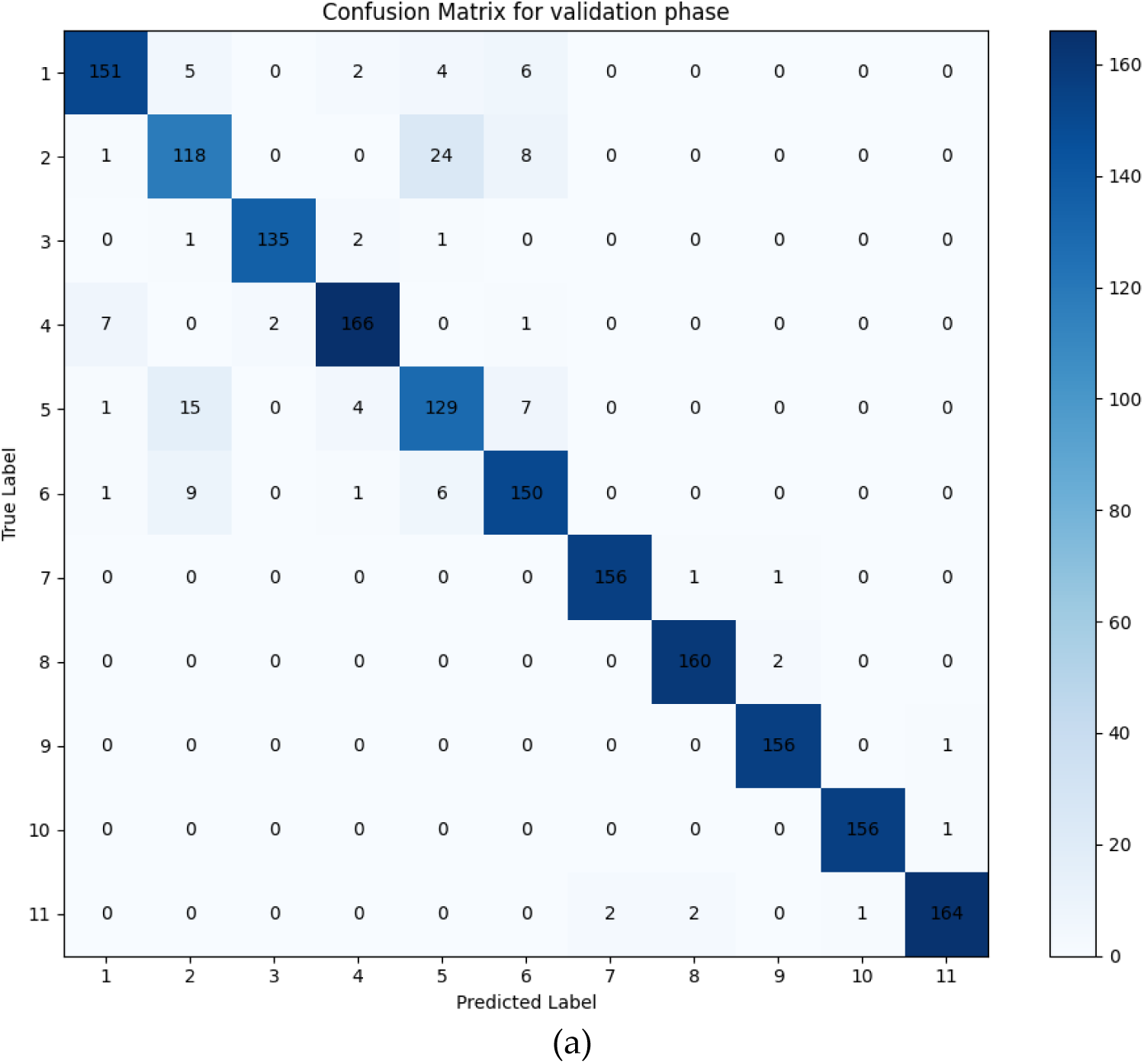
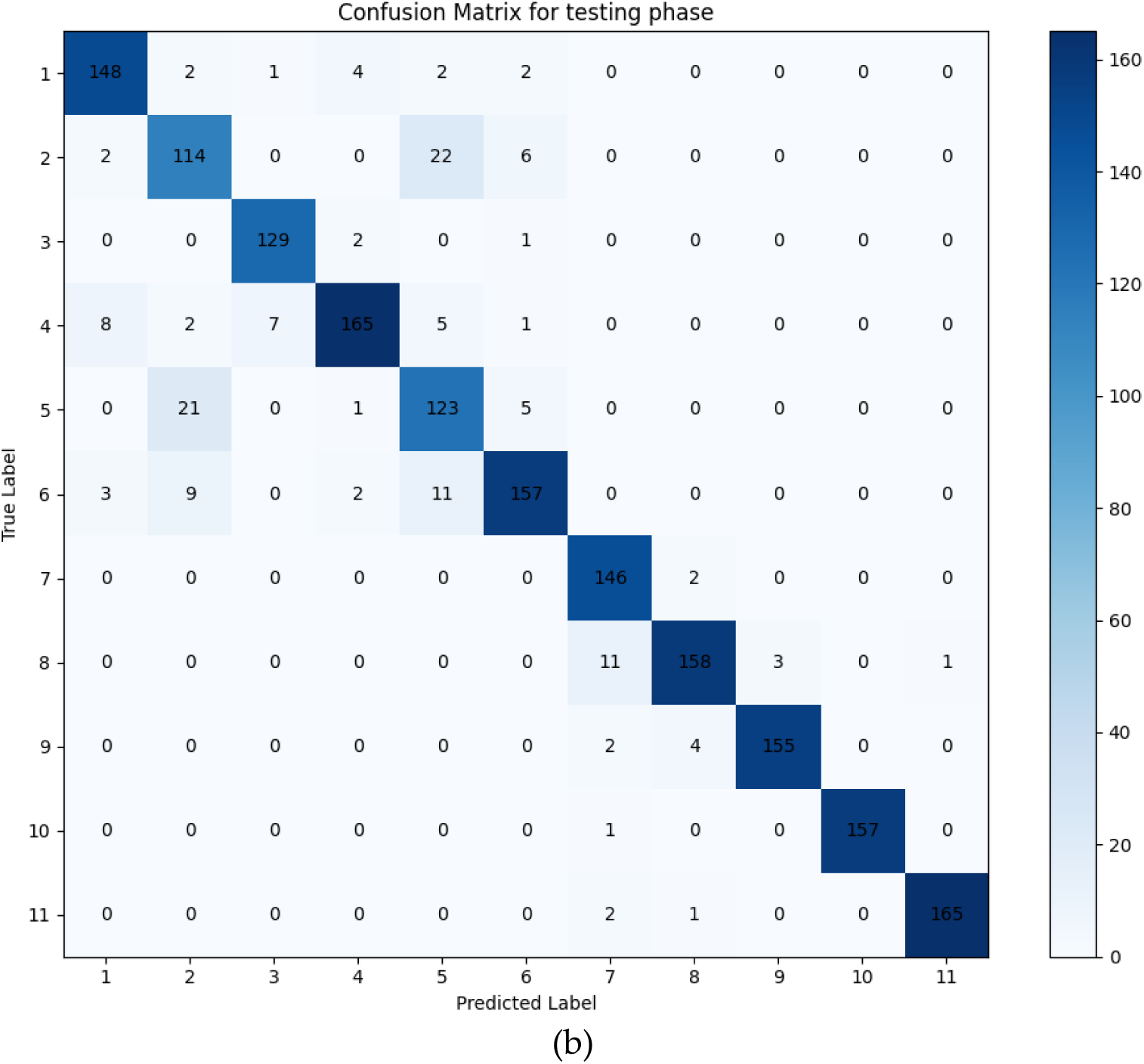
Confusion matrix (a) validation phase, (b) testing phase

In the testing phase confusion matrix, each row represents the true label (or class), while each column corresponds to the predicted label. The matrix displays the counts of instances where a certain true label was predicted as another label. For instance, the first row indicates that there were 148 instances of true label 1, out of which 148 were correctly predicted as label 1, 2 were predicted as label 2, 1 was predicted as label 3, and so forth. Similarly, for the second row, there were 114 instances of true label 2, out of which 114 were correctly predicted as label 2, 22 were predicted as label 5, 6 were predicted as label 6, and so on. This pattern continues for all true labels, providing insight into the model’s performance in the testing phase. The eleven categories include a wide range of anomalies, with the initial, third, fourth, seventh, and eighth groups being the ones that particularly describe aberrant kinds. The technique that we have developed indicates a considerable improvement in the precision of classification for these abnormal categories. With the significant ramifications, particularly in situations when prompt action is essential, such as the diagnosis of cancer, it is of the utmost importance that the model does not make the mistake of incorrectly identifying patients. Therefore, the efficiency of the model that we have proposed can be directly evaluated based on its capacity to accurately categorise aberrant kinds, which is a reflection of the possible influence that it may have on outcomes in the actual world.

**Table 7.**
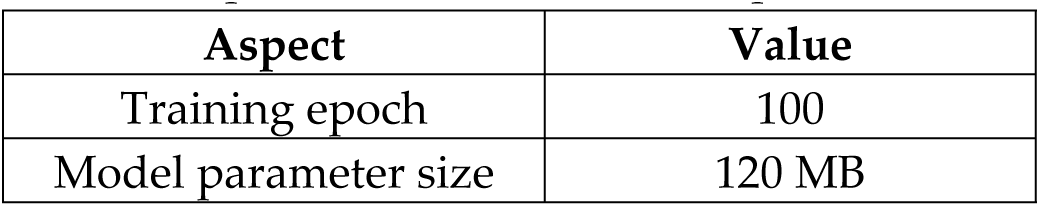
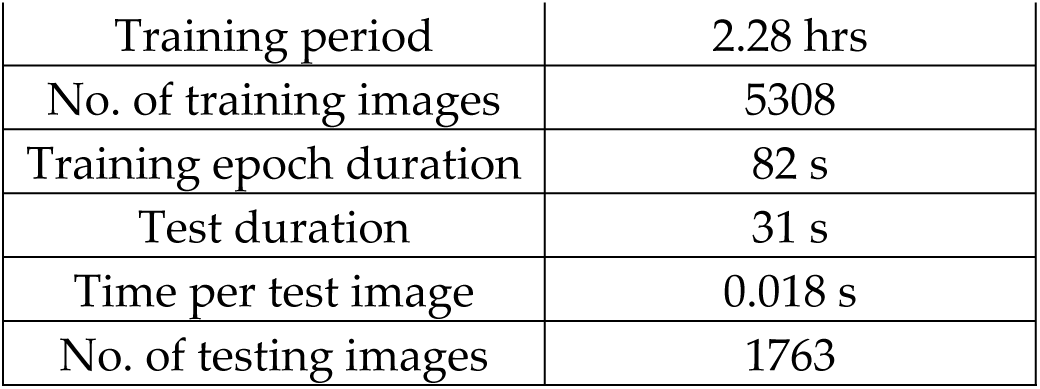
Proposed model utilized computational time.

Table 7 illustrates the suggested network model is trained in this experiment over 100 epochs, or around 82 seconds, every epoch. The total size of the model’s parameters is around 120 MB. To maximise the model’s performance, 5308 training images are used in total throughout the training phase. Even while the training phase takes a long time—roughly 1.3 hours—the testing phase that follows is much shorter—it lasts for only 31 seconds. A total of 1763 images are analysed during testing, and it takes an average of 0.018 seconds to analyse each test picture. When taken as a whole, these metrics provide valuable information on the time and computing resources needed for the model’s training and testing stages. Recently, a number of research have been carried out to obtain high classification accuracy in medical image analysis using different models and datasets. Naif Al Mudawi (2022) obtained an accuracy of 98.91% by using the UCI repository using SVM and XGBoost models. Shenghua Cheng (2021) employed a ConvNet model to achieve 98.12% accuracy with an emphasis on mother and child health. Hua Chen (2021) obtained a noteworthy accuracy of 99.08% by using a VGG model and the SIPaKMeD dataset. Using the CRIC dataset, Hiam Alquran (2022) used an SVM model and obtained a 97.3% accuracy rate. With MobileNetv2-YOLOv3 on the SIPaKMeD dataset, Lidiya Wubshet (2022) achieved 96.84% accuracy. Vijayanand (2021) obtained a 99% accuracy rate using ConvNet and DTCWT models on the Herlev dataset. Using a variety of models, including AlexNet, ResNet-18, ResNet-50, and GoogLeNet, Madhura Kalbhor (2023) obtained a 95.14% accuracy on the SIPaKMeD dataset. Using ConvNet, Javier Civit-Masot (2023) obtained a 97.1% accuracy rate on the Mendeley liquid cytology dataset. Using the SIPaKMeD dataset, Abinaya (2024) used a 3DCNN model and obtained a 98.6% accuracy rate. Sher Lyn Tan (2024) obtained a 99.2% accuracy rate using DenseNet-201 on the MDE-Lab dataset. Using the Risk Factors dataset from the ML repository, Jesse Jeremiah (2022) used LASSO and decision trees (DT) to achieve an accuracy of 98.72%. Lastly, the model that was suggested in 2024 made use of the SIPaKMeD and CRIC datasets, using SQ-ResNet152 to achieve an astounding accuracy of 99.74%. Together, these papers illustrate how medical image processing has advanced significantly and highlight the efficacy of different models on a range of datasets.

**Table 8.** Accuracy comparison of proposed and other models.

| Model | Accuracy (%) |
| --- | --- |
| SQ-ResNet152 | 99.74 |
| ResNet50 | 98.65 |
| VGG16 | 97.85 |
| DenseNet121 | 98.45 |
| InceptionV3 | 97.95 |
| MobileNetV2 | 97.30 |

The purpose of this study was to look into the several deep learning models that have been developed for cervical cancer classification and the enhancement of their accuracy when analyzing this disease shown in Table 8. The SQ-ResNet152 model was found to be the most accurate at 99.74%, showcasing the efficiency of its enhanced feature recalibration. A smaller form ResNet bundling labelled ResNet 50 was also up to the task recording an accuracy of 98.65%, this was then followed by DenseNet121 who topped at 98.45%. The VGG16 networks performed to 97.85% accuracy, even though they were unsuccessful in keeping up with modern architectural innovations like others, still had depth. InceptionV3 with complex inception modules achieved an accuracy of 97.95%. The light weight variant, MobileNetV2, which was made to ease workload, maintained the least accuracy at 97.30%, proving that there is a limitation on accuracies that can be obtained from such structures. These findings emphasize the different capabilities that various architectures possess in completing cervical cancer classification tasks.

**Table 9.** Classification accuracy comparison of proposed and other state-of-the-art methods.

| Author | Dataset | Model | Classification accuracy (%) |
| --- | --- | --- | --- |
| Hua Chen [13] | SIPaKMeD | VGG | 99.08 |
| Hiam Alquran [14] | CRIC | SVM | 97.30 |
| Lidiya Wubshet [15] | SIPaKMeD | MobileNetv2-YOLOv3 | 96.84 |
| Vijayanand [16] | SIPaKMeD, Herlev | DTCWT, ConvNet | 99.00 |
| Madhura Kalbhor [19] | SIPaKMeD | Alexnet, Resnet-18, Resnet-50 and Googlenet | 95.14 |
| Abinaya [22] | SIPaKMeD | 3DCNN | 98.60 |
| Proposed model | CRIC and SIPaKMeD | SQ-ResNet152 | 99.74 |

Table 9 illustrates the classification accuracy comparison of proposed and other state-of-the-art methods. The technique that has been created is able to distinguish 11 types of cervical cancer by analyzing the images of Pap smears utilizing the largest data collection ever put to this particular use. It remains a pioneer system in the field and is the fastest of them all. The ConvNet architecture that has been presented indicates a clearly higher classification rate as compared to past approaches. For the multi-class classification of pap smear images, recent machine learning techniques such as support vector machine (SVM), decision trees, K-means, and genetic algorithm have provided some insight into the possibilities. However, such models are limited by several constraints which include: a lack of effective cell clustering techniques when dealing with overlapping cells, overfitting with many factors, high-level intricacy which leads to time consuming, detection of rounded shapes only, poor accuracy, difficulty in optimizing parameters for nonlinear data sets, slowly trained models. The optimized SQ-ResNet152 model we have proposed and developed to solve these problems is also based on transfer learning. In the transfer learning-based optimized SQ-ResNet152 model that we have presented, we rather effectively perform the process of extracting important features from the pictures by virtue of enhancing classification performance. If we are to discuss the image classification tasks, in some cases, training deep neural networks from scratch is going to be a long and expensive affair in terms of computation. This challenge, our strategy is able to surmount through the application of transfer learning. We start off with a model that has already been trained on some knowledge of lower-level information. This is a cheap technique which increases the number of training samples required for convergence and reduces the time it takes for fine tuning. To make matters worse, the resources which are necessary to collect and annotate a large number of data sets for a particular task may not be practically feasible in many situations that exist in the real world. The transfer learning-based model that we have developed allows the usage of smaller datasets because the previously trained larger datasets are taken advantage of. Because of this even with a small quantity of labelled data, enhanced generalisation and performance is achieved. The accuracy comparison of proposed and other conventional models in Figure 12.

**Figure 12.**
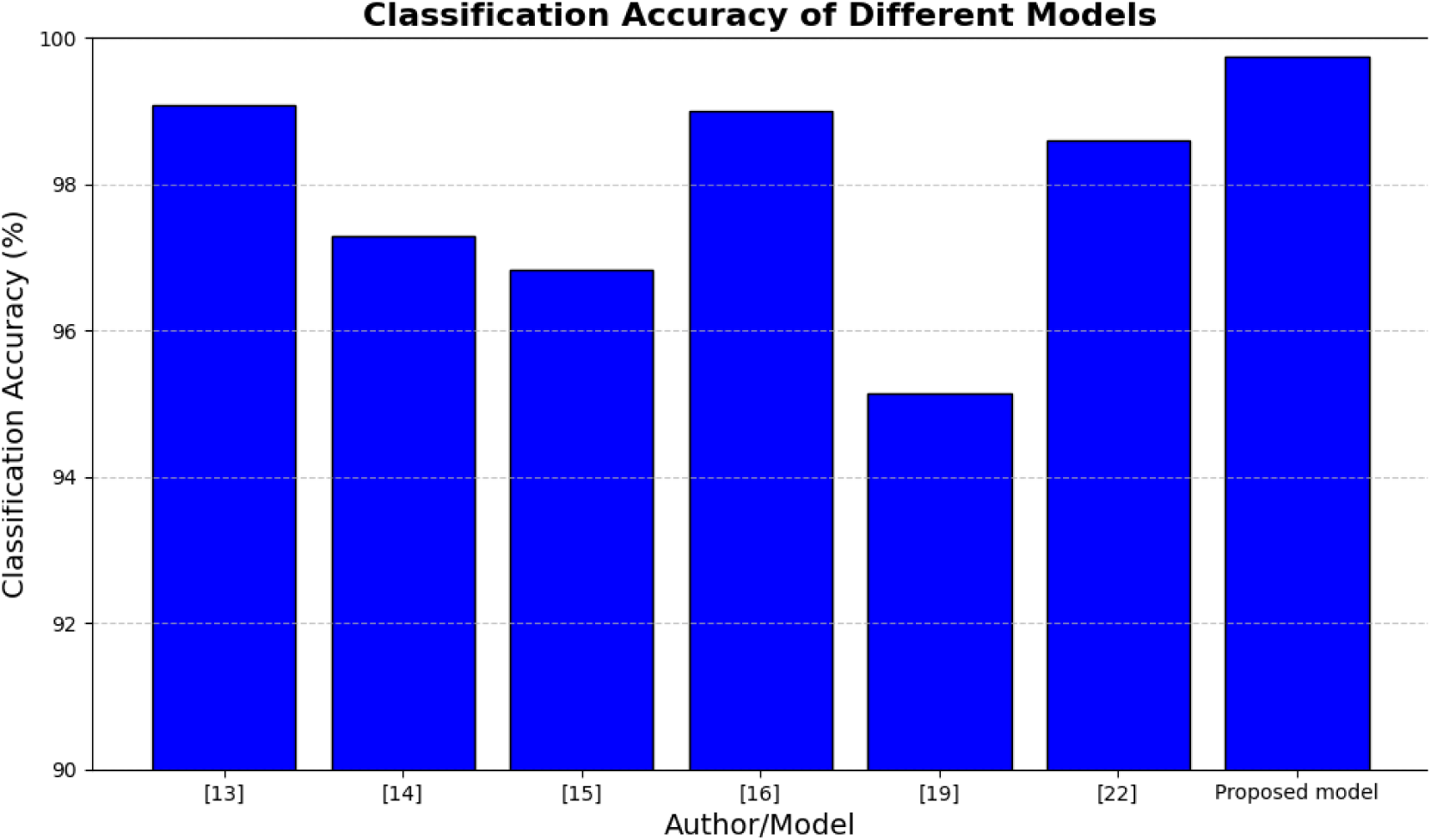
Classification accuracy comparison of proposed and other existing models

### 4.3 Ablation Study

This study presents all expected sequential phases of the work, such as the introduction, image dependent stages of deep learning model for cervical cancer classification in the framework of DHO and SE based modules effect. To start with, we analyzed the baseline performance of the ResNet152 model without the DHO optimization and found the standard levels of accuracy with increased training time for model convergence. That is why first we made a comparison with some of the standard optimization methods including Stochastic Gradient Descent (SGD) techniques and the DHO approach. The outcomes suggested that DHO was found to be more efficient in the tuning of the hyperparameters since it achieved fast convergence and better classification metrics. In the same manner, we applied ablation on the SE modules by first re-training the model without use of any SE blocks and in subsequent experiments incorporated use of SE modules in all layers of the network. The results of this study demonstrated that the addition of SE modules considerably increased the ability of the model to adjust the weights of important features resulting in a significant increase in accuracy and precision. However, upon application of SE modules to the last few layers of the model, we were able to achieve a nice equilibrium between computational cost and performance of the model. Taking into account all these problems, one can conclude that both DHO optimization and SE modules are of great importance to the improvement of medical image classification accuracy using deep learning models approaches.

## 5. Conclusion

The purpose of this research is to provide a unique deep learning framework that is specifically designed for cervical cell classification tasks. This framework is the transfer learning-based optimised SQ-ResNet152 model. The model is put through rigorous training and testing methods, which include the use of a massive collection of images. The first stages in the preprocessing process entail removing noise from the images and increasing their contrast quality. In order to categorise numerous cervical cancer illnesses across 11 different categories, the SQ-ResNet152 model, which is based on transfer learning, is then used in conjunction with a DHO optimizer to do the analysis of the picture dataset. Pap smear pictures may be distinguished with remarkable accuracy, sensitivity, and specificity according to the suggested model, which also greatly reduces the amount of time required for detection and the number of epochs necessary. The suggested transfer learning-based optimised SQ-ResNet152 model is capable of accurately classifying Pap smear pictures, achieving an amazing total accuracy of 99.74%. The results of the experiment on segmentation benchmarks demonstrate that the model is superior to numerous other deep-learning network models that are already in existence. It achieves performance that is equivalent to that of classic cervical cancer classification methods.

For the purpose of further optimising the technique that has been presented, we want to investigate a variety of additional model combinations in the future. Among these features are the enhancement of the capabilities for feature extraction and the refinement of the module structure. In addition, we want to study various data pre-processing methods, such as random cropping and colour jittering, in order to improve the generalisation capabilities of the model. The total effectiveness and performance of the method will be improved using these tactics, which will contribute to the improvement.

## Author Contributions

All authors contributed equally in this study. All authors have read this work.

## Institutional Review Board Statement

Not applicable.

## Informed Consent Statement

Not applicable.

## Data Availability Statement

This article uses only cropped images (4789 total) that display cervical cells [26]. The data used in this study is available on Kaggle dataset https://www.kaggle.com/datasets/prahladmehandiratta/cervical-cancer-largest-dataset-sipakmed.

## Acknowledgement

Not applicable.

